# SAME-clustering: Single-cell Aggregated Clustering via Mixture Model Ensemble

**DOI:** 10.1101/645820

**Authors:** Ruth Huh, Yuchen Yang, Yuchao Jiang, Yin Shen, Yun Li

**Author notes:** These authors contributed equally to this work (Ruth Huh, Yuchen Yang). Ruth Huh, Department of Biostatistics, University of North Carolina at Chapel Hill, Chapel Hill, NC 27599, USA.

## Abstract

Clustering is an essential step in the analysis of single cell RNA-seq (scRNA-seq) data to shed light on tissue complexity including the number of cell types and transcriptomic signatures of each cell type. Due to its importance, novel methods have been developed recently for this purpose. However, different approaches generate varying estimates regarding the number of clusters and the single-cell level cluster assignments. This type of unsupervised clustering is challenging and it is often times hard to gauge which method to use because none of the existing methods outperform others across all scenarios. We present SAME-clustering, a mixture model-based approach that takes clustering solutions from multiple methods and selects a maximally diverse subset to produce an improved ensemble solution. We tested SAME-clustering across 15 scRNA-seq datasets generated by different platforms, with number of clusters varying from 3 to 15, and number of single cells from 49 to 32,695. Results show that our SAME-clustering ensemble method yields enhanced clustering, in terms of both cluster assignments and number of clusters. The mixture model ensemble clustering is not limited to clustering scRNA-seq data and may be useful to a wide range of clustering applications.

## INTRODUCTION

Recent technological advances in single-cell RNA sequencing (scRNA-seq) have allowed researchers to catalog the transcriptomes across a large number of individual cells, empowering us to systematically study the heterogeneity at the cellular level. scRNA-seq has transformed the paradigm of genomic studies by investigating biology down to the single-cell resolution, which unveils information masked from the commonly used bulk RNA sequencing (RNA-seq). scRNA-seq analysis has led to, among others, the identification of existing and novel cell types, characterization of cells, prediction of cell fate, classification of tumor subpopulations, and investigation of cellular heterogeneity (1–3). Single cell clustering is a crucialstep to achieve above-mentioned utilities(4). For example, only after clustering the single cells, the following analyses can be meaningfully and conveniently carried out: identification and examination of cell type specific gene expression signatures, adjustment of cell type compositions for differential expression, and deconvolution of bulk RNA-seq expression data. Due to its importance, it is not surprising to find many existing scRNA-seq clustering methods (5–10). Unforunately, we find that clustering results from different methods are rather dissimilar (Supplementary Figure 1), which is consistent with literature (11,12) and not surprising because different methods employ different strategies for essential components of clustering (including choice of distance metric, dimension reduction, clustering approach and estimation of number of clusters) (Supplementary Table 1). Each scRNA-seq clustering approach has its own strengths and limitations. Thus, the use of two or more clustering methods is recommended for more accurate and comprehensive overview of cell clustering. However, when true (“gold-standard”) cluster labels are not available, it is difficult to select the best method(s), either before or after clustering analysis.

To address the challenging issue of selecting the optimal method(s) when true cell types are unknown, combining information from multiple individual methods becomes an appealing alternative. We present <u>S</u>ingle-cell <u>A</u>ggregated <u>C</u>lustering via <u>M</u>ixture Model <u>E</u>nsemble clustering (SAME-clustering), a well-grounded statistical model to solve the problem of consensus clustering. We use a cluster ensemble method because it is known to provide robust and improved quality solutions (13). Moreover, the multinomial mixture model cluster ensemble approach underlying SAME-clustering accommodates varying numbers of clusters from individual solutions, addresses the issue regarding correspondence of cluster labels across different solutions, and solves the issue of missing labels from some solution(s) (13,14). Furthermore, mixture model is a maximum likelihood-based approach where we can conveniently leverage model selection criterion to determine the optimal number of clusters for the final ensemble solution.

## MATERIAL AND METHODS

### Overview of SAME-clustering

In the current implementation of SAME-clustering, we first input a gene expression matrix into five individual clustering methods, SC3 (5), CIDR (6), Seurat (8), t-SNE (15)+ *k-means*, and SIMLR (9), to obtain five sets of clustering solutions. Of the five sets of solutions, we choose a maximally diverse subset of four according to variation in pairwise adjusted Rand index (ARI). The four individual solutions selected are then combined into a *n* × 4 matrix, where *n* represents the number of single cells. Inferred cell labels from the four sets of solutions are then used as input for the essential ensemble module of SAME-clustering, which assumes that these labels are drawn from a mixture of multivariate multinomial distributions (14) to build an ensemble solution by solving a maximum likelihood problem using the expectation-maximization (EM) algorithm. Figure 1 shows the overview of our SAME-clustering method.

**Figure 1.**
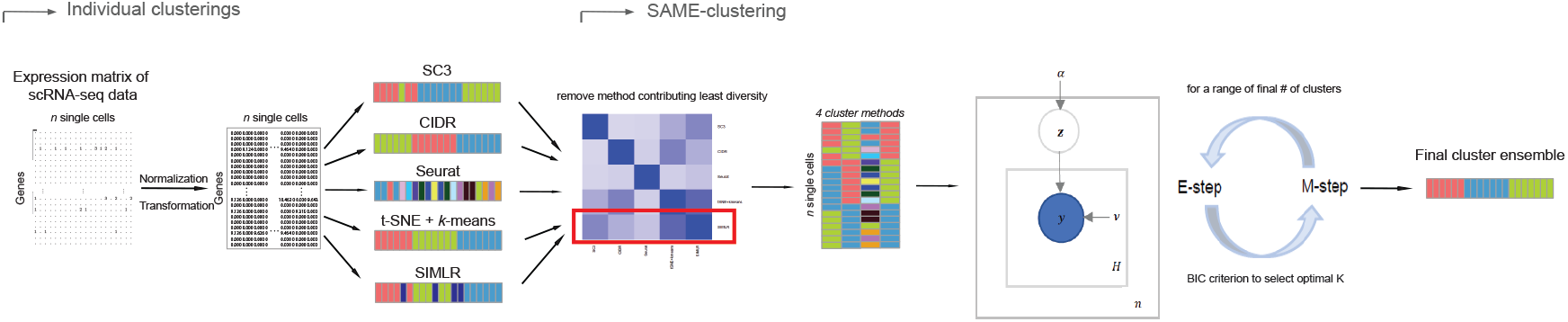
Overview of SAME-clustering. Normalization and transformation of scRNA-seq data are executed as specified by SC3, CIDR, Seurat, t-SNE+*k-*means, and SIMLR. From the 5 sets of clustering solutions, we take a subset of 4 diverse sets of clustering solutions. Then the *n* × 4 matrix, where *n* represents the number of single cells, is combined using SAME-clustering. Maximum likelihood estimation is achieved through EM algorithm and the optimal number of clusters is determined according to the BIC criterion, to provide the final consensus clustering solution. Notations are detailed in main text.

### Benchmark datasets

We assembled a total of 15 published datasets (1,3,7,16–22) that have “gold-standard” (deemed as true) cluster labels assigned to each single cell. For the Li dataset (7), there were seven cell lines, two of which have two batches per cell line. Since the individual clustering methods used in SAME-clustering, except for Seurat, do not have an option for batch effect correction, we kept only the larger batch for each of the two cell lines with two batches. We created our two large datasets by mixing single cells from purified peripheral blood mononuclear cells (PBMC) generated by 10X genomics (21), following Sun et al (10). Specifically, the simple case consists of three highly distinct cell types: CD56+ natural killer cells, CD19+ B cells and CD4+/CD25+ regulatory T Cells. In contrast, single cells from three similar cell types, namely CD8+/CD45RA+ naïve cytotoxic T cells, CD4+/CD45RA+/CD25- naïve T cells and CD4+/CD25 regulatory T cells, are combined to form the challenging case dataset. Supplementary table 2 shows the wide variety of published datasets that we adopted and tested SAME-clustering on.

### Implementation of the five individual clustering methods

SC3 filters out genes/ transcripts that are expressed in <10% or >90% of cells to remove rare and ubiquitous genes/transcripts. We applied SC3 with this default gene filtering step to all datasets to reduce dimensionality and without substantial impact on clustering results (Supplementary Table 3). For the simple and challenging PBMC mixture datasets, SC3 estimated 906 and 943 clusters respectively. As the number of clusters determined by the default method in SC3 performs unsatisfactorily for large datasets, we run principal component analysis (PCA) on the 1000 most variable genes, based on log2 transformed counts per million (CPM) data, to visually estimate the number of clusters. PCA plot shows three clear clusters for the simple case and two vague clusters for the challenging case (Supplementary Figure 2). We therefore used these numbers when applying SC3 to these PBMC mixture datasets.

For Seurat, dimension was reduced according to the number of principal coordinates (PCo) determined by CIDR implementation because CIDR provides an automatic way to select the number of PCos and SAME-clustering performs robustly across different number of PCs selected (Supplementary Figure 3, Supplementary Table 4). When applying t-SNE+*k-*means, we added an intermediate step to automatically detect the number of clusters and cluster centroids using ADPclust (23), which we have found previously to stabilize the performance of t-SNE+*k-*means(12). Although this automatic detection of centroids may affect cluster results for t-SNE+*k-*means when compared with manual inspection of centroids (Supplementary Figure 4 & 5, Supplementary Table 5), SAME ensemble results remained stable (Supplementary Table 5). For SIMLR, we set the number of clusters at the true values for the simple and challenging datasets, because the computational costs are high for SIMLR to determine the number of clusters for such large datasets.

Unlike SC3 and Seurat, CIDR, t-SNE+*k-*means, and SIMLR do not perform gene filtering. In order to test the potential impact of gene filtering on clustering performance, we adopted the method used by SC3. Specifically, we assessed the impact of different levels (ranging 0-10%) of filtering on individual methods (Supplementary Table 6) and the extent to which gene filtering affects SAME ensemble clustering (Supplementary Table 7). Our results suggest that gene filtering tends to improve clustering and we therefore conducted all our analysis with the 10% filtering criterion. More details on the implementation of these individual methods can be found in the Supplementary Data.

### SAME-clustering method

We adopted the notation laid out by Topchy (14). Suppose that we have *n* single cell data points *X* = {*x*_1_, …, *x*_*n*_}. *X* serves as input to various (five in SAME by default) individual clustering methods to return a set of labels. Let *H* be the number of individual clustering methods (by default, *H* = 4 as SAME selects four out of five), then for each data point *x*_*i*_ there would be *H* elements in the set of inferred cluster labels.

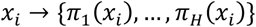

*π*_*j*_(*x*_*i*_) is the label assigned to single cell data point *x*_*i*_ by the *j*^*th*^ clustering method with *i* = 1,2, …, *n* and *j* = 1,2, … , *H*. We make no assumption for the correspondence of labels across the *H* methods. Hereafter, we will use the notation *y*_*ij*_ = *π*_*j*_(*x*_*i*_) or *y*_*i*_ = *π*(*x*_*i*_), which is the dataset used to generate our final consensus partition *π*_*C*_. This problem can now be considered as a categorical clustering problem, where we find a partition *π*_*C*_ of a set of vectors *Y* = {*y*_*i*_}.

We define

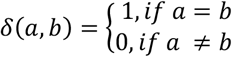

Then we model *y*_*i*_′*s* as random variables from a mixture of *M* multivariate densities. Each component *m* is parameterized by *θ*_*m*_, where *M* is the resulting number of clusters in the final consensus clustering and *m* = 1,2, …, *M*. *α*_*m*_ is the prior probability of the *m*^*th*^ cluster. The model assumes that *y*_*i*_′*s* are generated from first drawing from a cluster according to probabilities *α*_*m*_′*s*, then sampling a point from the cluster-specific density *P*_*m*_(*y*_*i*_|*θ*_*m*_). Taken together, we have:

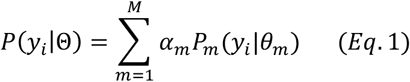

Assuming i.i.d for *y*_*i*_′*s*, the log likelihood function for {*α*_1_, …, *α*_*m*_, *θ*_1_, … , *θ*_*m*_) can be written as:

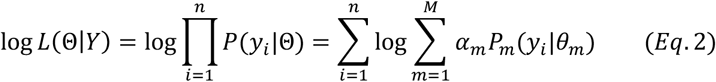

And now to find the best fitting mixture density for data *Y*, we need to maximize this likelihood function.

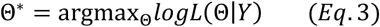

To simplify the problem, we assume conditional independence among individual clustering methods with each method weighted equally, so the conditional probability of *y*_*i*_ can be represented as the following:

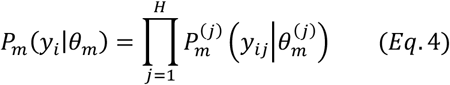

Since *y*_*ij*_ are nominal values reflecting cluster labels from partition *π*_*j*_, the probability density can be represented as a multinomial distribution:

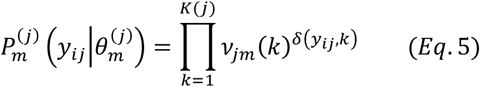

In (Eq. 5), *K*(*j*) is the number of clusters from partition *π*_*j*_, clustering result from the *j*^*th*^ method. The cluster label *k* in *π*_*j*_ therefore takes values from {1, …, *K*(*j*)}, and the probabilities of cluster labels are defined as 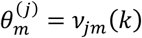, with the constraint that these probabilities sum to 1.

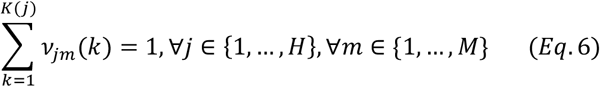

For a concrete example, when individual clustering method SC3 produced three clusters, (Eq. 5) can be represented as:

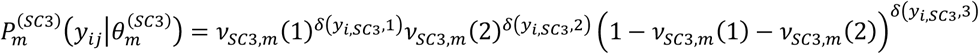

Note that each component *m* has a corresponding set of *v*′*s*, ∀*m* ∈ {1, …, *M*}.

One cannot analytically solve the maximum likelihood function in *Eq*. 3, when all the parameters (*α*’s and *v*’s) are unknown. Fortunately, however, we can optimize *Eq*. 1 via the EM algorithm. Specifically, we introduce hidden data *Z,* the distribution of which should be consistent with the observed values Y:

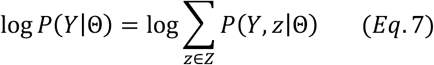

The values *z*_*i*_′*s* represent the mixture components used to generate data points *y*_*i*_′*s*. *z*_*i*_ = {*z*_*i*1_, …, *z*_*iM*_}, such that *z*_*im*_ = 1 if *y*_*i*_ belongs to the *m* -th component and *z*_*im*_ = 0, otherwise. The complete data (*Y*, *Z*) likelihood can be written as:

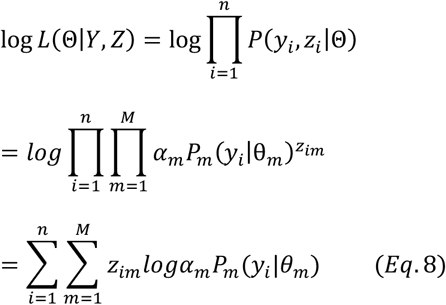

Following the general EM approach, the following function is defined, serving as the lower bound of the observed data likelihood:

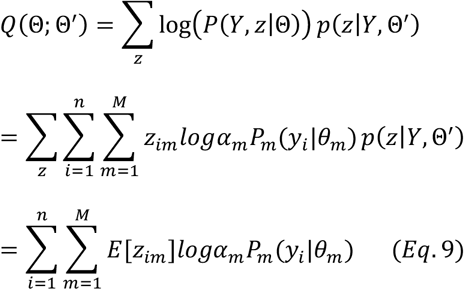

Maximizing *Q*(Θ; Θ′) is equivalent to maximizing the observed likelihood function in *Eq*. 2 where Θ = {*α*_1_, …, *α*_*m*_, *θ*_1_, … , *θ*_*m*_} (24,25). We optimize *Q* via the following Expectation (E) and Maximization (M) steps. In the E step, we compute *E*[*z*_*im*_] with current estimates of the parameters. In the M step, we apply the Lagrange multiplier along with the constraint ∑_*m*_ *α*_*m*_ = 1, and ∑_*k*_ *v*_*jm*_(*k*) = 1 to estimate the parameters with updated expected values of the hidden variables.

Expectation (E) Step:

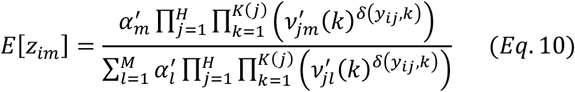

Maximization (M) Step:

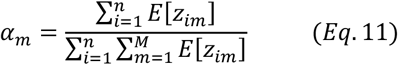

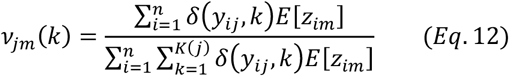

Each run of SAME initializes *α*′*s* and *v*′*s* according to a Dirichlet prior with a vector of 1’s. We repeat the E and M steps until convergence criterion is met. The convergence criterion we used for our implementation is *loglikelihood*^*Iteration*^ − *loglikelihood*^*Iteration*−1^ < 0.0001. After convergence is achieved, *y*_*i*_ is assigned to the component that has the largest expected value for the hidden label *z*_*i*_ = {*z*_*i*1_, …, *z*_*im*_}, as *E*[*z*_*im*_] denotes the probability that *y*_*i*_ is drawn from the *m*^th^ mixture component. This results in our final consensus partition *π*_*C*_. Since EM Algorithm does not guarantee reaching the global maximum (26,27), we run three chains of EM with random initializations by default in SAME as we observe ARI results stabilize with three chains of EM (Supplementary Figure 6).

### EM with missing labels

Some clustering methods, such as Seurat, give missing cluster labels for some of the data points. Therefore, our EM algorithm needs to be modified to accommodate such missing data. Now 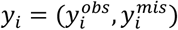. For the E-step, *E*[*z*_*i*_] is computed over the observed *y*_*i*_′*s*, denoted as 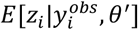, which means the product changes from 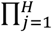 to 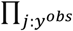. For the M-step, complete data log likelihood function is maximized to obtain estimates for parameters *v*_*jm*_(*k*) with 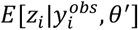 and 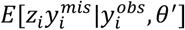 calculated and substituted into the M-step *Eq*. 11 and 12 (28). For missing observations in *Eq*. 12, *δ*(*y*_*ij*_, *k*) would be replaced by the current estimate of *v*_*jm*_(*k*).

### Determining the final estimated number of clusters

To determine the final number of clusters/ components, we run the multinomial mixture ensemble method multiple times over a range of *M* values. In our implementation, we run our ensemble SAME-clustering method for 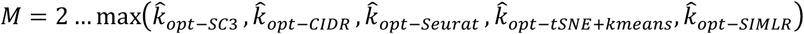. We calculate MLE by maximizing the following log likelihood function until convergence criterion is met.

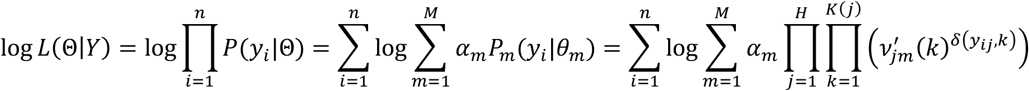

Next, we calculate AIC or BIC for each *M* we attempt the ensemble method, where *n* is the number of single cells, *p* is the number of parameters and 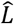 is the maximized value of the likelihood function.

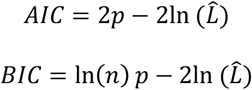

For the number of parameters, we have *M* − 1 parameters for the mixture weights, plus 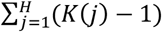 for each of the *M* component, leading to the total 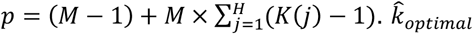 would be the *M* value that gives the lowest BIC or lowest AIC, depending on our choice of the model selection criterion. We found that BIC either outperforms AIC, or is merely slightly inferior to AIC in terms of ARI for the benchmark datasets (Supplementary Figures 7-9). Therefore, we choose BIC as our default model selection criterion.

### Diversity of individual cluster results to improve SAME-clustering

Existing literature has pointed out the importance of diversity in partitions from individual methods to enhance the performance of ensemble solution (29–32). To assess the diversity of the five individual clustering methods, we calculate pairwise ARI’s to quantify similarity between any two individual clustering solutions. Note that a low pairwise similarity represents a higher diversity. After attaining all the pairwise similarities, which can be represented as a heatmap, we calculate the variance for the vector of similarities with each method, including the similarity of value 1 for the method to itself. Due to the inclusion of this value 1 (self-similarity), this method-specific variance-based statistic tends to be larger for methods that are dissimilar to others. In comparison, the method with the lowest variance is most similar to other methods, with evidence aggregated from all pairwise comparisons. We, therefore, removed the method with the lowest variance, since the method would contribute the least in terms of diversity. We observed that this diversity-filtering approach results in improved performance in six datasets (Darmanis, Deng, Li, Baron_human1, Baron_human2, and Baron_human 3), same performance in two datasets (Ting and Yan), and impaired performance in five datasets (Supplementary Figure 10). The average increase in ARI for the six datasets (0.048) is slightly higher than the average decrease in the five datasets (0.038). Taking the reduced computational costs also into consideration, we proceeded with removing the method that contributes the least diversity.

## RESULTS

We benchmarked our SAME-clustering method and the five individual methods on 15 published datasets that represent a wide variety of sequencing technologies, tissue of origins, data units, numbers of single cells and numbers of cell types (Supplementary Table 2). Figure 2 summarizes clustering results, gauged by ARI. Among the 15 attempted datasets, SAME-clustering produces the best results in eight datasets (Darmanis, Baron_human1, Baron_human3, Baron_human4, Baron_mouse1, Goolam, Zeisel, and the challenging case), and the second best in four datasets (Biase, Baron_human2, Li, Yan). Additionally, SAME-clustering outperforms at least three individual methods in all 15 datasets. To further support consistency of SAME in producing reliable results we rank each method from 1^st^ to 6^th^ for all datasets, where ties are replaced by their mean rank. Figure 3 clearly shows that SAME-clustering outperforms all other methods rank-wise. The worst rank of 5.5 came from the Biase dataset, where only one cell was misplaced by SAME, leading to a high ARI but didn’t perform well rank-wise because there were 3 methods that achieved perfect clustering when compared to the “gold-standard”. We also compare our results to our previously published SAFE method (12), which performed overall second best and remains an attractive alternative (33), particularly when analyzing large datasets to save computation time.

**Figure 2.**
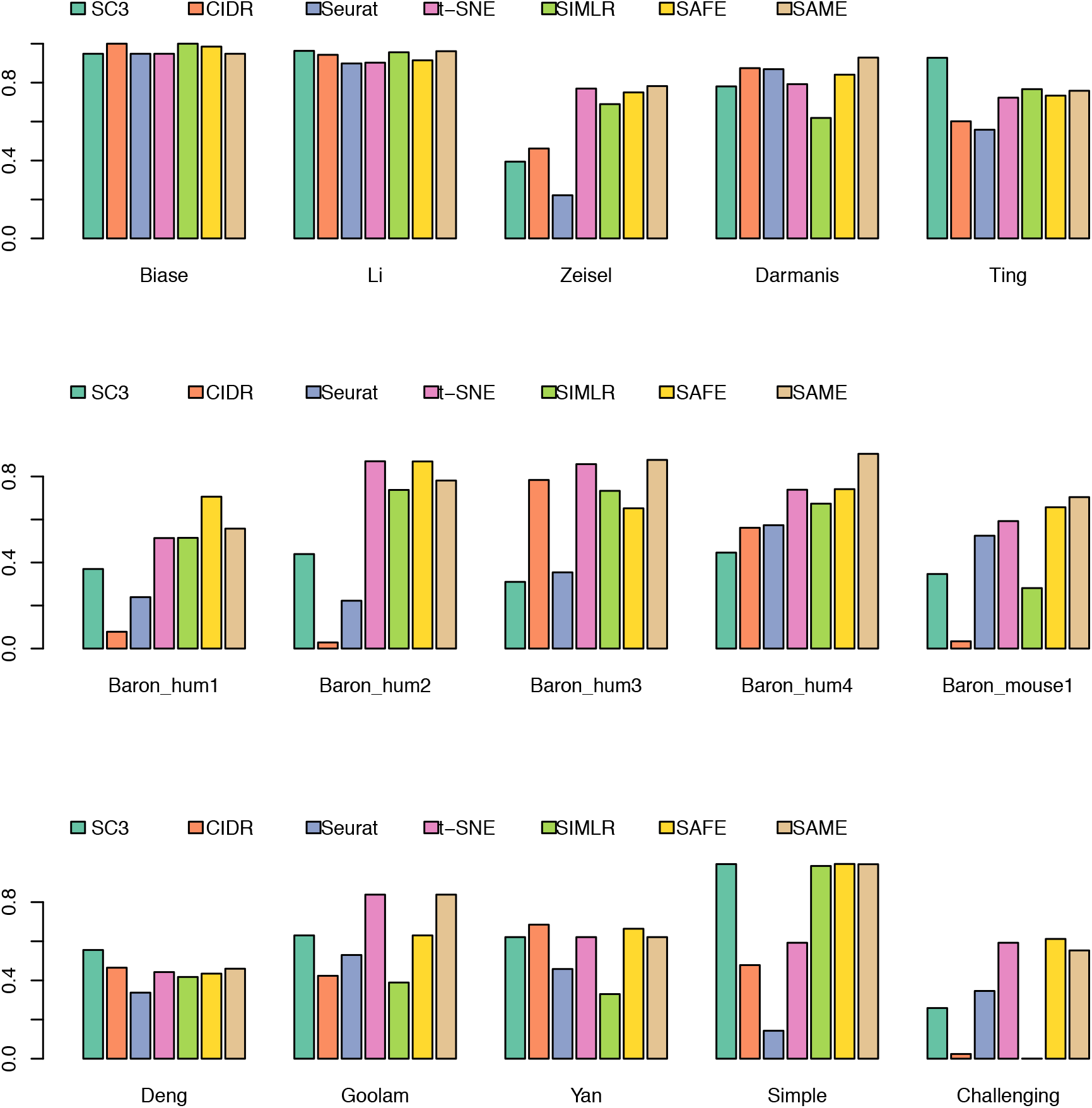
Benchmarking of SAME-clustering in fifteen published datasets. Similarity between estimated and “gold-standard” cluster labels is measured through Adjusted Rand Index (ARI), for 15 benchmark datasets.

**Figure 3.**
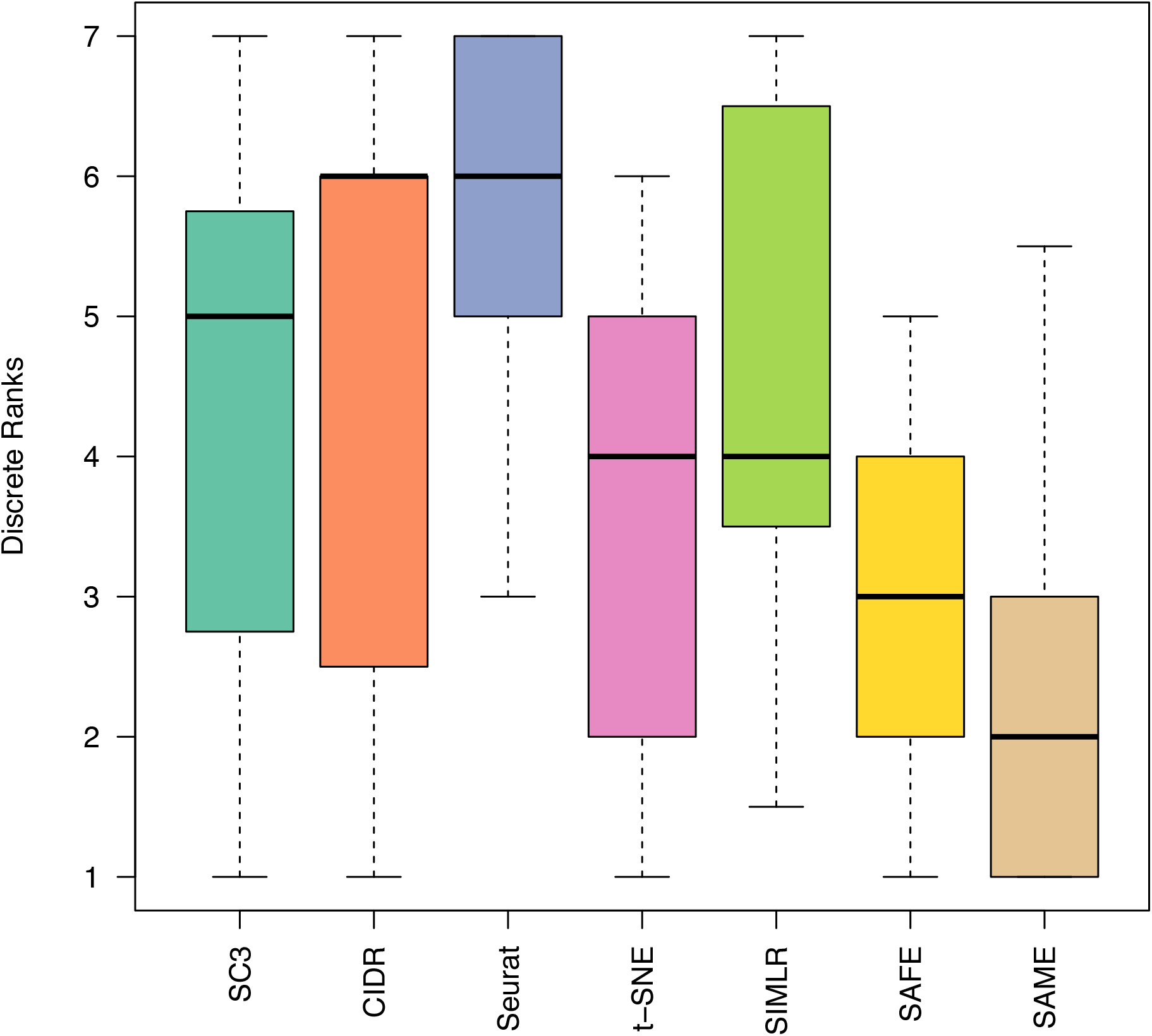
Method ranking across 15 datasets. Each method is ranked according ARI for 15 datasets. Lower rank represents better performance (1 is the best and 6 is the worst). Ties are replaced by the mean of their ranks.

We further investigated the performance of SAME-clustering in terms of estimating the number of clusters, compared to the five individual methods. Generally speaking, SC3 tends to overestimate the number of clusters, while CIDR, t-SNE+*k-*means, and Seurat underestimate the number of clusters. SIMLR performs reasonably well in most datasets achieving the best estimates. SAME-clustering also tends to underestimate the number of clusters, especially for the Baron datasets where the true number of clusters are either 13 or 14 (Supplementary Figure 11). For the five Baron datasets, there are five to eight rare cell types (defined as <1% of total single cells, with actual number of single cells ranging from 1-36). Individual clustering methods and SAME-clustering are not able to classify rare clusters accurately, which is a common problem with clustering methods. When we focus on the non-rare cell types, it becomes more apparent that SAME-clustering outperforms all other methods (Figure 4). Additionally, we have also assessed the several other computing and performance aspects of SAME-clustering, including the potential factors that may influence SAME’s performance, computing time and performance for large datasets, and simulation evaluation. Details are given in the Supplementary Materials (Supplementary Figures 12-23).

**Figure 4.**
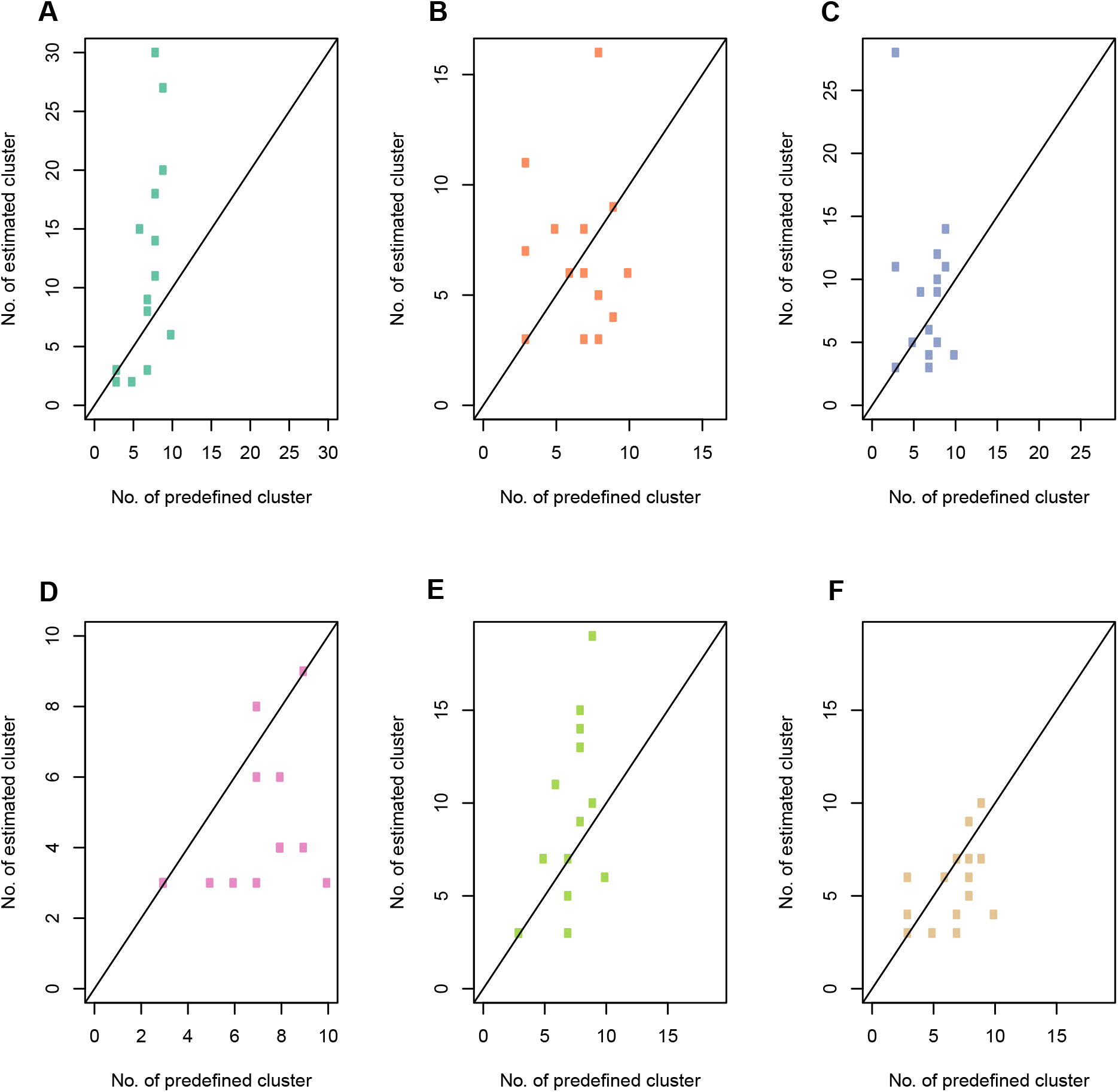
Accuracy of estimated number of clusters for non-rare cell types. Correlations between estimated and true number of clusters across 15 benchmark datasets, for SC3 (A), CIDR (B), Seurat (C), t-SNE+*k-*means (D), SIMLR (E), and SAME-clustering (F) respectively. Rare (<1% of total single cells) cell types are removed.

Interestingly, SAME-clustering demonstrates its capability to identify novel clusters. Figures 5 and 6 show an example from the simple case PBMC dataset, where SAME-clustering separates a novel cluster of single cells. According to the “gold-standard” labels defined by cell surface markers used in the original FACS experiments (34), there are three clusters in this dataset: CD56+ natural killer (NK) cells, CD19+ B cells and CD4+/CD25+ regulatory T Cells. The new fourth cluster is comprised of only 72 cells out of a total of 27,733 cells. Of these 72, 67 are *a priori* annotated as NK cells, 1 as a B cell, and 4 as regulatory T cells. *NKG7* and *GNLY*, which are known NK cell marker genes (21), are highly expressed in NK cells (as expected) but not in the new cluster of cells identified by SAME (Figure 5). Although most cells in the novel cluster are annotated as NK cells, the low expression of the known marker genes suggests that these cells have transcriptomic profiles deviating from the “gold-standard” NK cells. The remaining three clusters correspond to B cells, T cells, and NK cells. We feed cluster labels from SAME into *Seurat* to find the top ten cell type marker genes that are expressed in at least 70% of cells of the corresponding cell type. Figure 6 shows that, when sampling cells that have concordant labelling between the *a priori* annotation and SAME clustering, there is an apparent separation of B, T, and NK cells (Figure 6: left panel). In contrast, when we examine the expression profiles of cells in the new cluster, they separate into three groups. The first and third sub-groups (Figure 6: right panel: left most and right most columns) express marker genes of both regulatory T and NK cell types, and the second sub-group of single cells (Figure 6: right panel: middle columns) do not clearly express marker genes for any of the three cell types. These findings suggest that single cells in this new cluster, most (67/72) of which had “gold-standard” NK cell labeling from the original publication(21), are different from the typical NK cells and may represent intermediate cell type(s) that warrants further investigation.

**Figure 5.**
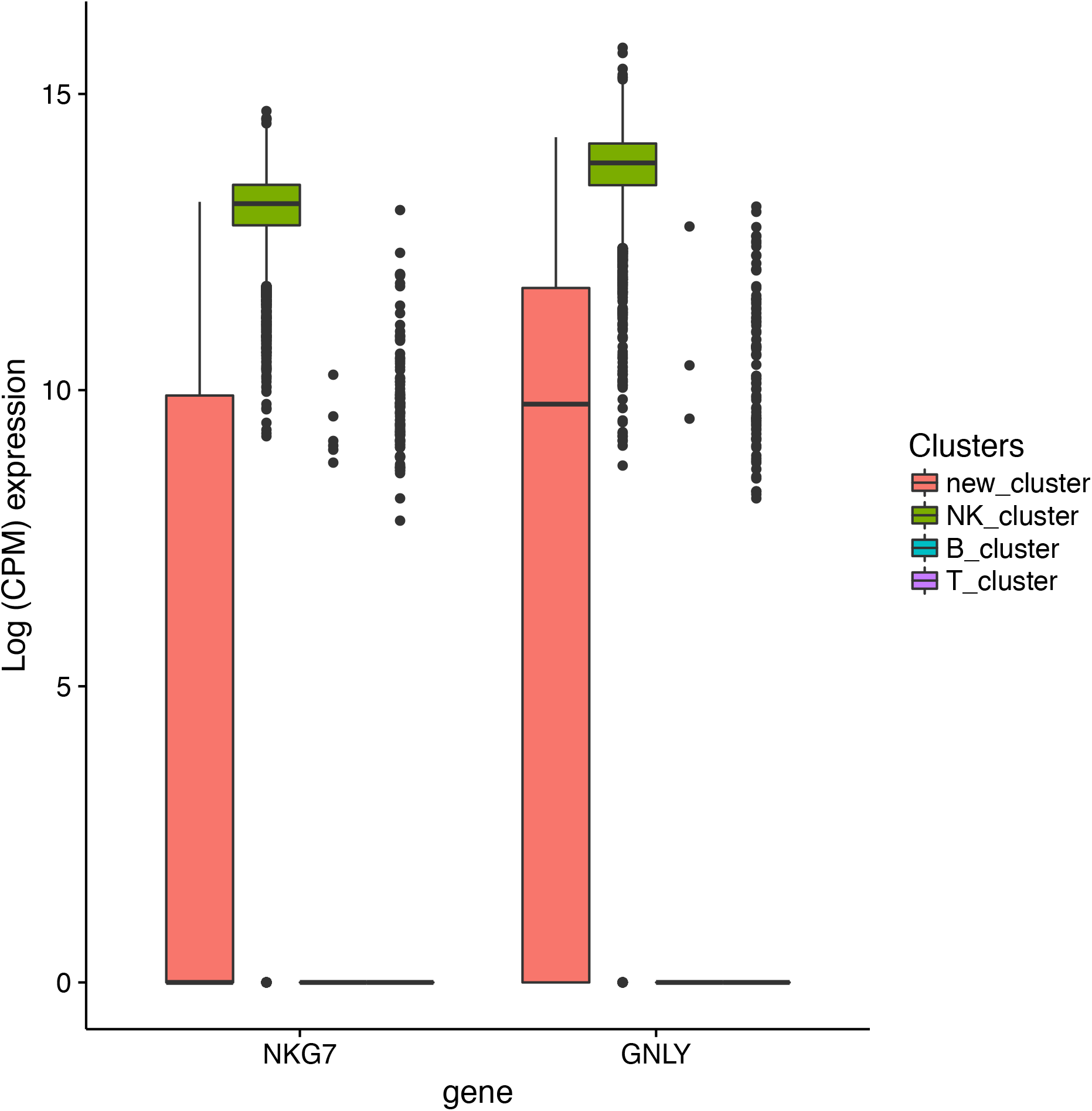
Identification of a novel cluster in the simple PBMC dataset. Expression of two known NK cell-type marker genes in cells labeled as NK cells by both “gold-standard: annotation and SAME-clustering (NK_cluster) vs cells in the novel cluster (new_cluster).

**Figure 6.**
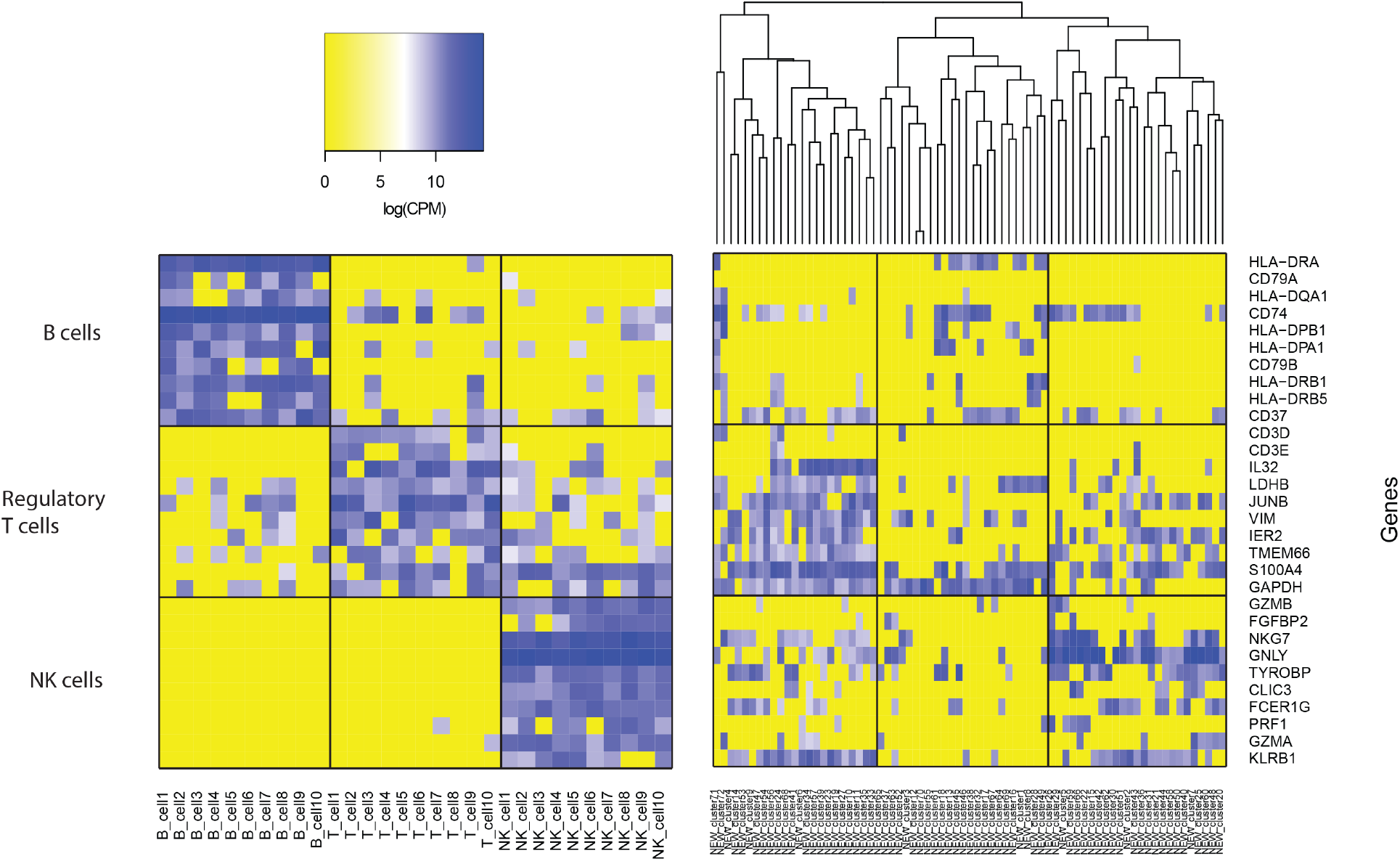
Marker gene expression for 30 selected single cells with concordant cell labels and 72 single cells in the novel cluster (simple PBMC dataset). Expression of marker genes specific to B, regulatory T, and NK cells. Left: expression heatmap of 30 single cells for which SAME cluster labeling agrees with gold-standard annotations. Right: heatmap of 72 single cells belonging to the new cluster identified by SAME.

Additionally, our SAME-clustering discovers limitations in “gold-standard” cell type annotations. We will illustrate with one example in the Darmanis (18) dataset. We combined top 40 cell-type markers of astrocytes, oligodendrocytes, and neurons from an independent study (35) and intersected them with *Seurat* to identify marker genes for each corresponding ensemble cluster, resulting in 24 astrocyte markers, 15 oligodendrocyte markers, and 7 neuron markers. Figure 7 (rightmost three columns) shows three discordant cells: one was annotated as an astrocyte but was identified as an oligodendrocyte by SAME (“true_astro_ens_oligo” in Figure 7); one was annotated as a neuron but was identified as an astrocyte by SAME (“true_neuron_ens_astro” in Figure 7); and one as annotated as an oligodendrocyte but was identified as a neuron by SAME (“true_olig_ens_neuron” in Figure 7). The first two discordant cells express marker genes from both their annotated cell type and the SAME-classified cell type. Such transcriptomic profile indicates that these single cells, containing mRNAs from signature genes of multiple cell types, may derive from a transient state, a doublet, or present a novel cell type. The last cell, although annotated as an oligodendrocyte, predominantly expresses neuron markers, suggesting that the original annotation might be problematic. In contrast, three randomly selected single cells from each of the three cell types, whose cluster labels by SAME agree with “gold-standard” annotations, show rather clean expression of marker genes from one single cell type.

**Figure 7.**
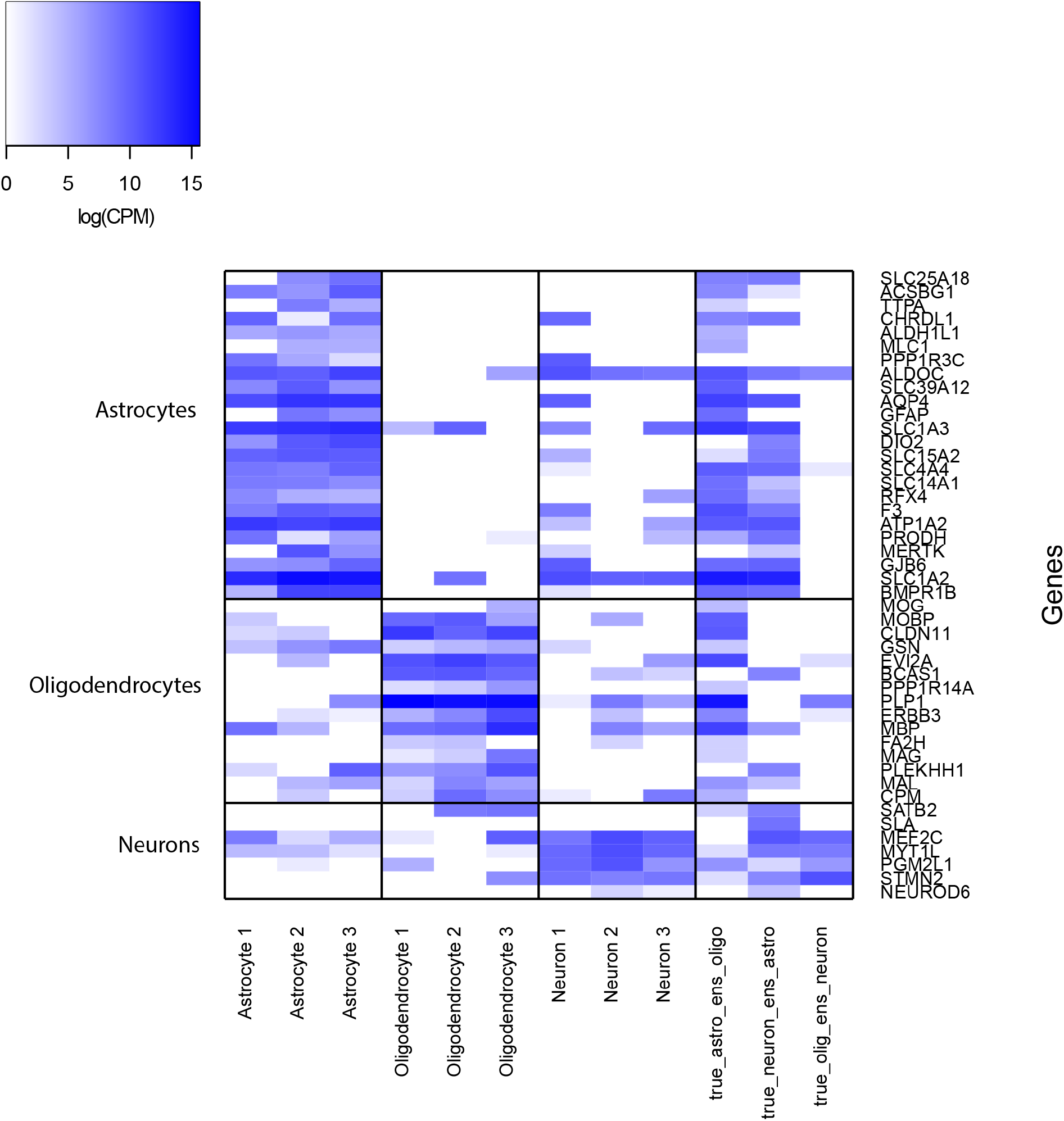
Revealing limitations in cell type annotations in the Darmanis Dataset. Expression of marker genes for Astrocytes, Oligodendrocytes, and Neurons. First 9 columns correspond to 9 single cells, by randomly selecting 3 cells from each cell type, for which SAME inferred labels agree with the “gold-standard” annotations. Last 3 columns correspond to single cells for which SAME disagree with the “gold-standard” annotations.

## DISCUSSION

We propose and implement SAME-clustering, a mixture model based probabilistic framework, that performs ensemble clustering for scRNA-seq data. Results across 15 real datasets show that SAME-clustering provides accurate and robust clustering. The ensemble method is flexible and users may choose to use different individual clustering methods. Not only does SAME-clustering provide improved clustering performance for labeling individual single cells, it also attains more accurate estimates for the number of clusters through the BIC model selection criterion. Overall, SAME-clustering offers an appealing solution to the scRNA-seq clustering problem, where individual clustering methods show substantial discrepancy (Supplementary Figure 1) with no method being an apparent winner across all datasets (Figure 2).

Ensemble modeling using mixture of multivariate multinomial distribution provides a well-grounded statistical framework that resolves the problem of lacking correspondence in labels across methods and naturally addresses the issue of missing cluster labels. SAME-clustering provides accurate, either the best or close match to the best, clustering results through combining diverse sets of clustering solutions obtained through varied dimensional reduction methods, distance metrics, and clustering methods. In addition, SAME-clustering provides more accurate estimate of the number of clusters compared to all individual methods examined through the well-established BIC statistical model selection criterion. Furthermore, SAME-clustering results are stable when parameters are changed for individual methods (Supplementary Tables 4 and 5). Our method is flexible and can easily accommodate additional sets of clustering solutions, as new clustering methods continue to be proposed (36). Supplementary Figure 10 shows that adding one more set of cluster results may improve ensemble results. We hypothesize that quality and added diversity of the additional contributing solution(s) influence whether the ultimate ensemble solution improves. However, these characteristics of individual cluster solutions are unknown *a priori*. Therefore, it is hard to gauge whether to include additional sets of cluster results. Our results (Supplementary Figure 10) suggest that diversity filtering of individual methods before ensemble leads to slightly improved ensemble clustering. The diversity filtering strategy is likely more useful when investigators choose to ensemble results from a larger number of individual methods, both in terms of clustering performance and computational costs. Our extensive evaluations demonstrate that SAME-clustering provides robust and accurate clusters for scRNA-seq data. Batch effect can potentially heavily influence clustering results. Although SAME has demonstrated satisfactory performances across real datasets, it is prudent and highly recommended to perform batch effect correction using customized batch effect correction methods (37) before running individual clustering methods and subsequently SAME. In addition, implementation of our SAME method provides a general and statistically rigorous framework for ensemble clustering using the mixture model-based method. We therefore expect SAME-clustering to be a helpful tool not only for single-cell clustering, but for other datasets that benefit from ensemble clustering approaches.

## AVAILABILITY

SAME-clustering, including source codes and tutorial, is available at https://yunliweb.its.unc.edu/same/ and https://github.com/yycunc/SAMEclustering.

