## Supplementary materials for "SAME-clustering: Single-cell Aggregated Clustering via Mixture Model Ensemble"

**S1. Individual methods capture different characteristics of scRNA-Seq data**

We show ARI (lower triangle) and NMI (upper triangle), both of which are similarity measures ranging from 0 to 1 with larger values corresponding to higher level of similarity, for every pair of clustering methods for the Zeisel dataset (1). ARI: Adjusted rand index; NMI: Normalized mutual information. ARI ranged (0.162-0.601) and NMI ranged (0.366-0.681). Numbers of clusters estimated by individual clustering methods are on the diagonals.

**S2. Implementation of the five individual clustering methods**

**S2.1. SC3**

SC3 includes the following steps. It first filters out genes/transcripts that are expressed in less than 10% of cells or more than 90% of cells to remove rare and ubiquitous genes/transcripts. 10% is the default value that can be adjusted by the user and this gene filtering step is optional as an entirety. After the filtering step, three distance metrics, namely Euclidian distance, Pearson and Spearman correlations, are calculated between single cells. These three distance matrices are then transformed using both PCA and eigenvector decomposition of a Laplacian graph separately. Then *k-*means clustering is performed on these transformed distance matrices. A consensus matrix of the different *k-*means clustering solutions is constructed using the cluster-based similarity partitioning algorithm (CSPA) (2). Lastly, this consensus matrix is clustered assuming *a priori* $\hat{k}_{opt\_SC3}$ clusters, where $\hat{k}_{opt\_SC3}$ is the SC3 estimated number of clusters. SC3 uses the Tracy-Widom distribution (3) to obtain $\hat{k}_{opt\_SC3}$. Although SC3 also applies cluster ensemble that aggregates over k-means clustering based on different distance metrics and dimensional reduction methods, the performance of SAME-clustering is more robust. It is mainly because SAME leverages information more broadly, over different individual clustering algorithms and includes more dimensional reduction methods (PcoA and t-SNE), distance metrics (learned similarity), and clustering methods (hierarchical clustering, modularity optimization).

SC3 employs a support vector machine (SVM) (4) model to speed up computation for large datasets. For datasets with more than 5,000 single cells, a subset of 5,000 single cells are randomly selected and clustered. The inferred labels are then used to train an SVM to assign labels to the remaining single cells.

SC3 reports that the gene filtering step does not affect clustering results but does reduce the dimensionality of the data and thus speeds up the algorithm. Our assessment of the effects of gene filter on the accuracy of clustering results corroborated their findings: Supplementary table 3 shows that gene filter indeed does not significantly affect clustering results. Adjusted rand index (ARI, ranging from 0 to 1 with higher values corresponding to higher similarity) (5) stays the same in four tested datasets with or without gene filtering. ARI improved with gene filtering in three other datasets and slightly decreased in the remaining six datasets. Supplementary table 3 also shows how the number of genes used (reflecting dimensionality), is reduced substantially after filtering, resulting in further reduced computational costs. Therefore, we applied SC3 with the default gene filtering step for all datasets.

For the simple and challenging PBMC mixture datasets, SC3 estimated 906 and 943 clusters respectively. A plausible reason is that large number of single cells together with low sequencing depth produced weak signals leading to suboptimal performance of the underlying algorithms (Kiselev, personal communication). Therefore, we performed PCA on the top 1000 most variable genes, based on log2 transformed CPM (counts per million reads) data, to visually estimate the number of clusters. PCA plot shows three clear clusters for the simple case and two vague clusters for the challenging case (Supplementary Figure 2). ARI for the simple case at *k*=3 is 0.994 and challenging case at *k*=2 is 0.259.

We tabulate ARI across datasets with and without the gene filter step. In the absence of gene filtering (i.e., Gene_filter = FALSE), all genes that are expressed in at least one single cell are counted. Bolded numbers mark datasets where gene filtering resulted in improved ARI. Italicized numbers mark datasets where gene filtering resulted in slightly decreased ARI.

**S2.2. CIDR**

CIDR’s main advantage is that it imputes dropout gene expression values to improve clustering performance. After dropout determination and imputation, dissimilarity matrix is calculated using squared Euclidean distance between pairs of cells. CIDR employs principal coordinate analysis (PCoA) for dimensionality reduction. For ease, CIDR *R* package automatically determines the number of principal coordinates to use for dimension reduction in its *nPC* function. However, users may choose to alter the number of PCo’s to use by searching for the elbow point in the plot showing the proportion of variation explained by each principal coordinate. With the dimension reduced dissimilarity matrix, hierarchical clustering is applied to assign cluster labels to single cells. The number of clusters is estimated with the Calinski-Harabasz index (6), denoted as $\hat{k}_{opt-CIDR}$.

**S2.3. Seurat**

Seurat combines dimension reduction with graph partitioning methods. The expression matrix is first filtered to exclude single cells with less than 200 expressed genes, and to exclude genes expressed in fewer than three single cells. Thus, Seurat will not give a final cluster label to single cells with < 200 expressed genes, resulting in missing cluster labels. For counts input, Seurat normalizes, for each single cell, by the total expression and multiplies by a scale factor of 10,000. Natural log transformation is subsequently applied after normalization. We skip the normalization step and only apply log transformation when input data are already normalized. Seurat then scales the data by removing unwanted sources of variation, by regressing out number of molecules detected per cell for UMI data or sum of non-normalized values within a cell for non-UMI data. Furthermore, Seurat can regress out batch effect, cell alignment rate, percent of mitochondrial genes, and cell cycle. Next, PCA is performed on the scaled data to reduce dimensionality. To cluster the cells, Seurat first constructs a *k-*nearest neighbor (KNN) graph with Euclidean distance in PCA space, where edges are drawn between similar cells. These edge weights are refined based on Jaccard distance which measures the dissimilarity between local neighborhoods. The cells are clustered by applying modularity optimization techniques (7), and the estimated number of clusters, $\hat{k}_{opt-Seurat}$, is determined during the process.

Among Seurat parameters, we have previously found that the resolution parameter is influential on results and decided on the value of 1.2 for datasets with less than 200 cells and the default value of 0.9 for larger datasets (8). To avoid manual inspection and determination of the number of PCs to use, we use the same number of PCs as PCos determined by CIDR, which as aforementioned provides an automatic way to select the number of PCos. Alternatively, users can examine the plot of the standard deviation of principal components and determine a threshold at the elbow of the plot. This decision, however, can be rather arbitrary when there is not a clear elbow (Supplementary Figure 3). Fortunately, we find that although using different numbers of dimensions may change the Seurat clustering results, it does not affect the consensus results as much. For four datasets, we evaluated the similarity of Seurat results to true labels using the number of PCs determined by CIDR (as part of SAME’s default automatic determination), and at several points that appear to be the “elbow” point (via manual/visual inspection). For the Biase and Darmanis datasets, the number of PCs selected by *nPC* function of CIDR, produced the best Seurat and SAME-clustering (Supplementary Table 4A-B). ARI of SAME-clustering ensemble solution in Biase dataset stayed the same (ARI of 1) across a range of number of PCs used, even when Seurat results varied substantially: ARI ranging for 0.344-0.948 (Supplementary Table 4A). Although ARI of SAME-clustering was not highest at the number of PCs selected by CIDR in Baron_human1, variation in ARI was lower in SAME-clustering compared to Seurat clustering in Biase, Darmanis, and Baron_human1. The higher variation of ARI for SAME-clustering in Baron_Human4 is due to a large improvement in SAME-clustering ARI using number of PC selected by CIDR. SAME-clustering always performed better than Seurat across a range of number of PCs selected (Supplementary Table 4A-D). Therefore, our results suggest that using the number PCs determined by CIDR’s number of PCo’s selected is reasonable for robust and high-quality Seurat clustering. More importantly, SAME-clustering results are robust to the choice of number of PCs used for Seurat clustering.

**S2.4. t-SNE+*k-*means**

t-SNE is a popular approach to visualize high dimensional data in two-dimensional space and has been widely used to visualize single cell clusters (9,10). After reduced to two-dimensional space, data can be clustered via *k-*means. The R package *Rtsne*, performs an initial reduction of space using PCA, and implements the Barnes-Hut algorithm (11) to mitigate the heavy computational burden that comes with minimizing the Kullback*-*Leibler divergence using gradient descent for large datasets. Default perplexity parameter of 30 is decreased to 10 for datasets with fewer than 200 single cells (8).

Both t-SNE and *k-*means are stochastic with varying clustering results across different initial points. To mitigate fluctuation across runs, we first apply ADPclust (12) to t-SNE transformed data to estimate the number of clusters as well as the cluster centroids. ADPclust is an adaptive density peak detection method using nonparametric multivariate kernel density estimation. To determine the number of clusters, $\hat{k}_{opt-tsne+kmeans}$, we evaluate cluster assignments through the silhouette index (13) using a grid of values for both bandwidth and number of clusters. ADPclust chooses a plug-in bandwidth estimator that minimizes the asymptotic mean integrated squared error (AMISE) to estimate the parameters of the multivariate kernel distribution. The cluster centroids and number of clusters that produces the maximum average silhouette index are carried forward to subsequent *k-*means clustering detailed below.

To test whether the number of clusters and initial centroids determined by ADPclust above lead to reasonable results for t-SNE+*k-*means clustering, we first visually inspected the cluster results. Biase and Li dataset showed clear separation in clusters, suggesting t-SNE+*k-*means generated satisfactory clustering results (Supplementary Figure 4A-B). For the three Baron datasets and Zeisel dataset, some of the smaller clusters were grouped into a bigger cluster. This is because ADPclust tends to choose a smaller number of clusters than what visual inspection suggests (Supplementary Figure 4C-F). We reran t-SNE+*k-*means by selecting centroids according to visual inspection, to assess the effects on ARI for both t-SNE+*k-*means clustering and SAME-clustering. Based on centroids manually selected according to visual inspection, Supplementary Figure 5 A-D show new cluster solutions of the same single cells in Supplementary Figure 4 C-F. Consistent with improved cluster separation based on these manually selected centroids, ARI of t-SNE+*k-*means improved in two of the four datasets (Supplementary Table 5 A-D). In terms of the final SAME-clustering ensemble results, selection of centroids via visual inspection showed no clear advantage: gauged by ARI, performance was improved in Baron_human2 and Baron_mouse1 but impaired in Baron_human4 and Zeisel datasets. In sum, manual determination of centroids from t-SNE visualization tends to improve t-SNE+*k-*means clustering when there is a clear global structure but can have negative effects when there is no clear global structure in the visualization (Supplementary Figure 5D) or when *k-*means’ tendency to produce equal-sized clusters leads to worse clustering results (Supplementary Figure 5B). Integrating information from other individual clustering methods, SAME is robust to how centroids are selected for *k-*means.

**S2.5. SIMLR**

Single-cell interpretation via multi-kernel learning (SIMLR) (14) takes raw gene expression matrix as input and applies a log10 transformation prior to analysis. SIMLR learns a similarity metric through learning proper weights for several Gaussian kernels. The advantage of using multiple kernels lies in its flexibility in comparison to a single kernel and therefore, can capture diverse statistical characteristics of single-cell data. Then a cell-to-cell similarity matrix in constructed accordingly and SIMLR assumes that this matrix should be approximately block*-*diagonal when clusters exist. This similarity matrix is used as input, instead of the regular gene expression matrix, to reduce the dimension of the data using t-SNE and *k-*means is applied subsequently for clustering.

For large datasets, SIMLR uses *k-*nearest-neighbor (KNN) to approximate the pairwise similarity matrix. After the similarity matrix is obtained, spectral clustering is adopted instead of applying *k-*means after t-SNE, because t-SNE is more computationally expensive. SIMLR supplies two ways to estimate the optimal number of clusters: eigengap and separation cost. Both utilize eigenvalues and eigenvectors which can easily become computationally expensive with larger datasets. Therefore, it is not feasible to estimate number of clusters using these methods for large datasets. Both measures select the same number of clusters in many cases. In our SAME implementation, we will use eigengap to estimate the optimal number of clusters for SIMLR,  $\hat{k}_{opt-SIMLR}$. Since the computational complexity is high for determining the number of clusters for large datasets, we set the number of clusters at their true values for the simple and challenging case.

**S3. Gene-Filtering for CIDR, t-SNE+*k-*means, and SIMLR**

Unlike SC3 and Seurat, which have a gene filtering step in the implemented software package, CIDR, t-SNE+*k-*means, and SIMLR do not have a gene filtering step. In order to test the potential impact of gene filtering on the performance of these three clustering methods, we adopt the method used by SC3. To remove lowly and highly expressed genes, SC3 filters out genes/ transcripts that are expressed in less than 10% of cells or more than 90% of cells. Similarly, we remove genes/transcripts expressed in less than X% of cells and more than (100-X)% of cells, where X takes values 0, 0.1, 1, 5, 10. From Supplementary Table 6, we reach no clear conclusion regarding the optimal level of gene filtering.

We further evaluated the effects of gene filtering by comparing the difference in ARIs of CIDR, t-SNE+*k-*means, SIMLR, and SAME-clustering, with and without a gene filtering step (with the default X=10%) (Supplementary Table 7). From the standard deviation of differences, we can see that CIDR (0.229) and SIMLR (0.182) vary rather substantially. t-SNE+*k-*means is not affected much by the gene filtering step. SAME-clustering shows more variation than t-SNE+*k-*means but is more robust compared to CIDR and SIMLR. SAME-clustering differences has a standard deviation of 0.127, and is driven by impaired performance in datasets that are not filtered. For example, without the gene filtering step, SAME-clustering’s ARI for Goolam is 0.32 less than when the dataset is filtered. Baron Human_4 also deteriorates substantially without the filtering step with ARI decreasing by 0.22. Considering the importance of gene filtering and to be more consistent with SC3 and Seurat, we carry on all our analysis with the 10% gene filter criterion.

**S4. Improving the EM Algorithm**

Since EM Algorithm does not guarantee reaching the global maximum (15,16), it is recommended practice to run multiple chains of EM with different initial points and retain the one with maximal likelihood. To guide our choice regarding the number of EM chains, we run SAME for five different datasets, setting the number of EM chains at 1, 2, 3, 5, and 10 and assess stability in the resulting ARI’s. From Supplementary Figure 6, we can see that there is either a large variation or extreme outliers in the five datasets when EM is run merely once or twice. Results then start to stabilize as variation decreases and the median stabilizes. We therefore run three chains of EM by default in SAME, consistent with the choice recommended in Topchy (17).

**S5. Empty Component(s)**

When running EM for a fixed *M* and a random initial point, some clusters (out of *M*) may end up empty (i.e., with zero single cells falling into the particular cluster[s]). To accommodate such scenario, we re-run EM up to three times until the number of non-empty clusters is equal to *M*, that is, no empty clusters. If the estimated non-empty number of clusters is still less than M after the third attempt, SAME clustering comes to an early stop, implying the number of clusters is less than the current M attempted.

**S6. AIC vs BIC**

AIC and BIC are both commonly adopted criteria for model selection. They both assess goodness of fit through the likelihood function, using different terms to penalize overfitting. In our context, AIC and BIC are both applied and evaluated to select the optimal number of clusters by selecting *M*, the number of components/clusters that minimizes the AIC or BIC criterion. We examine ARI's of thirteen datasets for both criteria. Supplementary Figure 7 shows that AIC and BIC result in the same ARI for four datasets (Biase, Darmanis, Li, and Yan); BIC outperforms in seven datasets (Baron_human1, Baron_human2, Baron_human3, Baron_human4 Baron_mouse1, Goolam, Zeisel); and AIC outperforms in two datasets (Deng and Ting). BIC either outperforms AIC, or is merely slightly inferior to AIC. Therefore, we choose BIC as our default model selection criterion.

Furthermore, BIC more accurately predicts the true number of clusters compare to the AIC criterion (Supplementary Figures 8 and 9). Additionally, as AIC creates more clusters, similar cells are divided into different clusters which causes less cluster similarity compared to BIC that groups similar cells together.

**S7. A concrete example for the number of parameters to be estimated**

| **Method** | **# of resulting clusters** | **Parameters** |
| --- | --- | --- |
| SC3 | 7 | $\nu_{1m}\left( 1 \right),\nu_{1m}\left( 2 \right), \nu_{1m}\left( 3 \right),\nu_{1m}\left( 4 \right),\nu_{1m}\left( 5 \right),\nu_{1},\left( 6 \right),\nu_{1m}\left( 7 \right)$ |
| CIDR | 4 | $\nu_{2m}\left( 1 \right),\nu_{2m}\left( 2 \right), \nu_{2m}\left( 3 \right),\nu_{2m}\left( 4 \right)$ |
| Seurat | 5 | $\nu_{3m}\left( 1 \right),\nu_{3m}\left( 2 \right), \nu_{3m}\left( 3 \right),\nu_{3m}\left( 4 \right),\nu_{3m}\left( 5 \right)$ |
| t-SNE+kmeans | 3 | $\nu_{4m}\left( 1 \right),\nu_{4m}\left( 2 \right), \nu_{4m}\left( 3 \right)$ |

The table above enumerates parameters that are required for the multinomial distribution to represent each individual clustering method, which depends on the number of clusters the individual method produces. Each component $m$has a corresponding set of $\nu's$, where $m=1,\ldots,M$. $M$is the number of clusters we want to produce with our ensemble method. Lastly, $\alpha_{1},\ldots,\alpha_{M},$ mixing coefficients are additional parameters that need to be estimated as well. These parameters are estimated through the EM algorithm with a set convergence criterion. After the convergence criterion is met, the expectation of the hidden variable $Z$ is used to assign an ensemble label to the single cell.

**S8. Computational costs**

To measure the computation time of SAME, we simulated four sets (mimicking four individual clustering methods) of clustering results, where single cells form 3 - 17 clusters, and performed SAME-clustering. We considered seven different numbers of single cells, ranging from 1,000 (1k) to 100,000 (100k). The SAME cluster ensemble step was repeated three times. The results show that, SAME needs ~0.05 - 10 hours for a dataset with 1k - 20k single cells (Supplementary Figure 12). When the number of single cells further increases, the running time varies substantially: ~21 - 66 hours for a dataset with 50k single cells, and ~22-124 hours for 100k single cells. The much larger variation in computing time for datasets with 50K and 100K single cells is partially due to larger variability in the performance of the computing nodes available on our research computing cluster when jobs require longer running time. Finally, we have also measured memory usage of SAME-clustering when applied to large datasets. We found < 1 GB memory RAM is required for a dataset with 100k single cells. Thus, for large datasets, there is no need to use any computation computing node with big memory for SAME-clustering.

**S9. Performance evaluation using simulated data**

We also performed the following simulations to evaluate the performance of our SAME-clustering. Specifically, we simulated 1,200 cells of three cell types with 15,000 genes following zero-inflated negative binomial distribution, and performed clustering using all the five individual clustering methods. The results show that, in this simulated dataset, CIDR works the best (ARI =1.00), followed by SC3 and t-SNE + k-means, while Seurat performs the worst (ARI = 0.002) (Supplementary Figure 13). Then we applied SAME-clustering both using all the five individual solutions and only four solutions without Seurat results. Under both two scenarios, SAME performs very well (ARI = 0.998 and 1.00 with or without Seurat results, respectively). The simulation results indicate the robustness and accuracy of SAME-clustering.

**S10. Additional performance evaluation**

In this study, we further evaluated several other computing and performance aspects of SAME-clustering. First, we assess SAME’s robustness to multiple runs of one original individual clustering algorithm. We applied t-SNE + k-means method to six datasets 100 times with different seeds, and then performed cluster ensemble using SAME-clustering. The results show that, in all the six datasets, SAME’s performance is comparable or better than multiple runs of t-SNE + k-means, as manifested by the ARI index (Supplementary Figure 14). These results suggest that, SAME-clustering not only can ensemble across different methods of large divergence, it can also handle cluster ensemble from multiple runs from the same individual algorithm.

Secondly, we evaluated SAME’s robustness to random clustering. Specifically, we simulated a set of random cluster labels for four datasets, and applied SAME-clustering to each dataset with this random set of clustering labels included. Each simulation setting was replicated 10 times with different random seeds. The results show that, as expected, the ARI of the randomly assigned cluster labels is close to 0, which means the clustering is indeed the expected random label assignment (Supplementary Figure 15). In contrast to the original version without the random cluster (SAME), there is no noticeable drop in clustering accuracy in all of the four datasets, which shows that SAME-clustering is robust and produces stable results even with the introduction of a random cluster.

To evaluate the influence of different numbers of input solutions on SAME’s performance, we have experimented SAME-clustering to use two to five maximally diverse sets of clustering results. The results show that, ensemble clustering using four diverse or all the five individual solutions perform the best in all the six tested datasets, while the results using fewer sets of individual clustering results, especially using only two diverse sets, are not satisfactory in most datasets (Supplementary Figure 16). These results suggest that incorporating four maximally diverse individual methods will enhance or at least stabilize the ensemble solution of SAME-clustering, which can also reduce the computational load (in contrast to using solutions from all individual methods), especially when there are more input individual solutions.

Besides the most commonly used ARI, there are several other indices, such as Rand Index (18) and silhouette width, that can be used to measure clustering accuracy. Here we applied both two indices to assess the performance of SAME-clustering. According to Rand Index, in the 12 benchmarking datasets, SAME-clustering outperforms all the individual solutions in five datasets, and performs better than at least two individual methods in another six datasets (Supplementary Figure 17). Similarly, the results of silhouette width show that, SAME-clustering performs better than all the individual methods in three out of the 12 datasets, and better than at least two individual methods in eight datasets (Supplementary Figure 18). These evaluations show that, SAME-clustering performs well across various datasets, using ARI, Rand Index and silhouette index.

To evaluate the influence of different gene sets used on clustering accuracy, we randomly sampled 50% of genes from the entire gene sets in four datasets and applied each of the five individual clustering methods to these datasets. Compared to the original results when the entire set of genes was employed, the clustering performance using a subset is comparable (Supplementary Figure 19).

We also compared the number of clusters estimated by SAME-clustering to the average number of clusters between different methods. The results show that, the cluster numbers estimated by SAME is closer to the ground truth (the average absolution deviation from the true cluster numbers is 1.933) than the average number across individual solutions (the average absolution deviation is 2.72) (Supplementary Figure 20). It indicates that SAME-clustering provides the most accurate estimation on the number of cell types compared to the individual methods.

In SAME-clustering, ADPclust was employed to detect the number of clusters and cluster centroids for t-SNE + kmeans clustering. Since ADPclust itself is a clustering method, here we also assess the clustering performance using ADPclust directly. Specifically, ADPclust was applied to the dimensional reduced data by t-SNE, and compared its performance to our original clustering using ADPclust + k-means. The results show that the performance of clustering using ADPclust only and ADPclust + k-means are rather comparable, except in Ting dataset (Supplementary Figure 21). We then performed SAME-clustering with ADPclust solutions, and also observed slightly worse results using ADPclust only than the original ADPclust + k-means. Thus, we would still keep the original implementation of including k-means.

To evaluate SAME’s performance in large and complex datasets, we applied SAME ensemble clustering on the “Fresh 68k PBMC dataset”, which consists of several T cell sub-clusters of high similarity, such as CD8+ cytotoxic, CD4+, memory and regulatory T cells (19). Because of the large number of cells, we only used three individual methods that handle such a large dataset, SC3, Seurat and SIMLR. For both SC3 and SIMLR, because estimating the number of clusters (k) is memory consuming, we performed both methods using two a priori selected k’s: k = 10 and 11. The two values were selected according to the numbers of clusters in the two sets of “truth” released from the original paper. Specifically, one set of the “true” labels is based on RNA-seq of purified PBMCs (here, k1-truth=11), and the other set of “truth” is generated by k-means clustering (here, k2-truth=10). We compared the ensemble results from SAME to the corresponding “truth” with the pre-selected k for SC3 and SIMLR. The results showed that, when using the first set of “truth”, SAME ARI is 0.411, outperforming all the three individual solutions (Supplementary Figure 22 right). When the second set is used as the “truth”, ARI of SAME is 0.554, excelling SC3 and SIMLR and slightly worse than Seurat (ARI=0.593) (Supplementary Figure 22 left). For the 100k simulated datasets, SAME was found to perform better than three out of the four individual solutions (Supplementary Figure 23). These results suggest that our SAME-clustering scales well and performs reasonably on large and complex datasets.

**S11. The novel PBMC cluster**

The novel cluster identified was not discovered by the majority of clustering methods. For this dataset, the four individual methods that were ensembled were SC3, Seurat, t-SNE+kmeans and SIMLR. Of the 4 individual methods, SC3, t-SNE+kmeans, and SIMLR classified the cells into 3 groups, whereas Seurat produced 28 clusters. After ensembling, SAME detected a new 4th cluster. As stated in the main text, “Of these 72, 67 are *a priori* annotated as NK cells, 1 as a B cell, and 4 as regulatory T cells”, where the *a priori* annotations (i.e., cell labels) were derived from FACS according to cell surface markers (19). For the 67 FACS sorted NK cells, we report below their cell type assignments by SC3, t-SNE+kmeans, and SIMLR. Of the 67, 22 are misclassified by SC3 (18 are grouped with B cells and 4 are grouped with T cells), 20 are misclassified by t-SNE+kmeans (18 are grouped with B cells and 2 are grouped with T cells), and 66 are misclassified by SIMLR (all 66 are grouped with T cells). 17 cells are misclassified by all three methods that detect 3 clusters. Again, Seurat results are much more complicated: these 67 cells fell into 9 out of the 28 clusters.

**Supplementary Table 1. Summary of characteristics of individual clustering methods.** Information includes existence of functionality for dropout gene imputation and filtering, choice of distance measures, dimension reduction, and clustering method usedand how cluster numbers are estimated.

| **Method** | **Dropout genes imputation** | **Filtering** | **Distance Metric(s)** | **Dimension Reduction** | **Clustering Method** | **Estimating # of clusters** |
| --- | --- | --- | --- | --- | --- | --- |
| SC3 | No | Remove rare and ubiquitous genes | Euclidean  Spearman  Pearson | PCA  Laplacian | *k*-means, hierarchical clustering | Tracy-Widom Distribution |
| CIDR | Yes | none | Euclidean | PCoA | Hierarchical clustering | Calinski-Harabasz index |
| Seurat | No | Remove rare genes and cells with rare expression | Euclidean | PCA | Modularity Optimization | Modularity Optimization |
| t-SNE+ k-means | No | None | Euclidean | PCA | K-means | ADPclust |
|  |  |  |  | t-SNE |  |  |
| SIMLR | No | None | learned similarity | t-SNE | K-means | eigengap |

**Supplementary Table 2. Major characteristics of the 15 benchmark datasets.** Information regarding organism, source, sequencing technology, measuring units of the gene expression matrix and number of single cells are included.

| **Dataset** | **Organism** | **Technology** | **Source** | **# of single cells** | **Units** | **Ref** |
| --- | --- | --- | --- | --- | --- | --- |
| Baron human 1 | Human | inDrop | Pancreas tissue | 1,937 | UMI | (20) |
| Baron human 2 | Human | inDrop | Pancreas tissue | 1,724 | UMI | (20) |
| Baron human 3 | Human | inDrop | Pancreas tissue | 3,605 | UMI | (20) |
| Baron human 4 | Human | inDrop | Pancreas tissue | 1,303 | UMI | (20) |
| Baron mouse 1 | Mouse | inDrop | Pancreas tissue | 822 | UMI | (20) |
| Biase | Mouse | SMART-Seq SMARTer | Embryo development | 49 | FPKM | (21) |
| Darmanis | Human | SMART-Seq SMARTer | Brain tissue | 420 | gene counts | (22) |
| Deng | Mouse | Smart-seq, Smart-seq2 | Embryo development | 286 | RPKM | (23) |
| Goolam | Mouse | Smart-seq2 | Embryo development | 124 | CPM | (24) |
| Li | Human | SMART-Seq SMARTer | Cell Line | 460 | FPKM | (25) |
| Ting | Mouse | scRNA-Seq, modified Tang2010 protocol(26) | Different tumors | 149 | CPM | (27,28) |
| Yan | Human | scRNA-Seq, modified Tang2010 protocol(26) | Embryo development | 90 | RPKM | (26,28,29) |
| Zeisel | Mouse | Quantitative scRNA-seq with UMI | Brain tissue | 3,005 | UMI | (1,30) |
| Simple | Human | 10X Genomics GemCode | PBMC | 28,733 | UMI | (10) |
| Challenging | Human | 10X Genomics GemCode | PBMC | 32,695 | UMI | (10) |

**Supplementary Table 3. SC3 performance (measured by ARI) across datasets, with or without gene filter.**

| Dataset | Gene_filter | | # of single cells | # of genes/transcripts | ARI |
| --- | --- | --- | --- | --- | --- |
| Biase | | TRUE | 49 | 13265 | 0.948 |
|  |  | FALSE | 49 | 25068 | 0.948 |
| *Darmanis* | | TRUE | 420 | 10743 | *0.728* |
|  |  | FALSE | 420 | 21517 | *0.788* |
| **Deng** | | TRUE | 286 | 10229 | **0.555** |
|  |  | FALSE | 286 | 21219 | **0.421** |
| Goolam | | TRUE | 124 | 11119 | 0.630 |
|  |  | FALSE | 124 | 28147 | 0.630 |
| Li | | TRUE | 460 | 20238 | 0.963 |
|  |  | FALSE | 460 | 43723 | 0.963 |
| **Ting** | | TRUE | 149 | 12764 | **0.833** |
|  |  | FALSE | 149 | 18594 | **0.795** |
| Yan | | TRUE | 90 | 6462 | 0.621 |
|  |  | FALSE | 90 | 14489 | 0.621 |
| *Zeisel* | | TRUE | 3005 | 9542 | *0.347* |
|  |  | FALSE | 3005 | 19970 | *0.359* |
| *Baron Human 1* | | TRUE | 1937 | 5835 | *0.339* |
|  |  | FALSE | 1937 | 16016 | *0.398* |
| **Baron Human 2** | | TRUE | 1724 | 5728 | **0.413** |
|  |  | FALSE | 1724 | 16310 | **0.409** |
| *Baron Human 3* | | TRUE | 3605 | 5167 | *0.307* |
|  |  | FALSE | 3605 | 16678 | *0.310* |
| *Baron Human 4* | | TRUE | 1303 | 6517 | *0.419* |
|  |  | FALSE | 1303 | 15720 | *0.426* |
| *Baron Mouse1* | | TRUE | 822 | 4725 | *0.324* |
|  |  | FALSE | 822 | 14359 | *0.343* |

**Supplementary Table 4. Robustness of Seurat and SAME-clustering across a range of #PCs selected**. A-D. ARIs for Biase (A), Darmanis (B), Baron_human1 (C) and Baron_human4 (D), respectively, across various numbers of PCs selected either by CIDR’s *nPC* function or by exploring the plot and standard deviation of PCs for a clear elbow. Standard deviation of ARIs for Seurat and SAME are calculated separately for robustness. Best performance across number of PCs used are bolded.

A Biase dataset

|  | # of PC selection method | | | | |  |
| --- | --- | --- | --- | --- | --- | --- |
|  | CIDR | Elbow plot | | | | sd |
| # of PCs | 2 | 4 | 5 | 6 | 7 |  |
| Seurat ARI | **0.948** | 0.733 | 0.744 | 0.411 | 0.344 | 0.252 |
| SAME-clustering ARI | **1** | **1** | **1** | **1** | **1** | 0 |

B Darmanis dataset

|  | # of PC selection method | | | |  |
| --- | --- | --- | --- | --- | --- |
|  | CIDR | Elbow plot | | | sd |
| # of PCs | 6 | 10 | 11 | 12 |  |
| Seurat ARI | **0.869** | 0.775 | 0.773 | 0.752 | 0.052 |
| SAME-clustering ARI | **0.914** | 0.882 | 0.882 | 0.879 | 0.017 |

C Baron_human1 dataset

|  | # of PC selection method | | | | | |  |
| --- | --- | --- | --- | --- | --- | --- | --- |
|  | CIDR | Elbow plot | | | | | sd |
| # of PCs | 4 | 8 | 9 | 10 | 11 | 12 |  |
| Seurat ARI | 0.239 | 0.432 | **0.546** | 0.513 | 0.538 | 0.540 | 0.120 |
| SAME-clustering ARI | 0.493 | 0.554 | 0.562 | 0.532 | **0.597** | 0.596 | 0.039 |

D Baron_human4 dataset

|  | # of PC selection method | | | |  |
| --- | --- | --- | --- | --- | --- |
|  | CIDR | Elbow plot | | | sd |
| # of PCs | 6 | 8 | 9 | 10 |  |
| Seurat ARI | 0.574 | 0.611 | 0.594 | **0.617** | 0.019 |
| SAME-clustering ARI | **0.918** | 0.701 | 0.682 | 0.626 | 0.128 |

**Supplementary Table 5. Impact of centroid detection strategy on clustering results.** Number of clusters k and ARI of t-SNE+*k-*means and SAME-clustering results, when centroids are selected either automatically by ADPclust or manually from t-SNE visualization. The datasets analyzed are Baron_human2, Baron_human4, Baron_mouse1, and Zeisel in A-D respectively.

A Baron_human2 dataset

|  |  | centroid detection strategy | |
| --- | --- | --- | --- |
|  |  | Automatic from ADP | Manual from visual inspection of t-SNE plot |
| tsne+*k-*means | k | 3 | 6 |
|  | ARI | 0.643 | 0.830 |
| SAME-clustering | k | 8 | 7 |
|  | ARI | 0.804 | 0.812 |

B Baron_human4 dataset

|  |  | centroid detection strategy | |
| --- | --- | --- | --- |
|  |  | Automatic from ADP | Manual from visual inspection of t-SNE plot |
| tsne+*k-*means | k | 3 | 6 |
|  | ARI | 0.732 | 0.629 |
| SAME-clustering | k | 9 | 8 |
|  | ARI | 0.705 | 0.697 |

C Baron_mouse1 dataset

|  |  | centroid detection strategy | |
| --- | --- | --- | --- |
|  |  | Automatic from ADP | Manual from visual inspection of t-SNE plot |
| tsne+*k-*means | k | 4 | 6 |
|  | ARI | 0.589 | 0.669 |
| SAME-clustering | k | 5 | 6 |
|  | ARI | 0.711 | 0.742 |

D Zeisel dataset

|  |  | centroid detection strategy | |
| --- | --- | --- | --- |
|  |  | Automatic from ADP | Manual from visual inspection of t-SNE plot |
| tsne+*k-*means | k | 4 | 7 |
|  | ARI | 0.769 | 0.672 |
| SAME-clustering | k | 10 | 11 |
|  | ARI | 0.661 | 0.576 |

**Supplementary Table 6.** **Impact of gene filtering on clustering performance of CIDR, t-SNE+*k-*means, and SIMLR.** To remove rare and ubiquitous genes/transcripts, we remove those expressed in less than X% or more than (100-X)% of cells. We test a range of X values from 0-10%. Highest ARIs are bolded.

|  | ARI for CIDR clustering | | | | |
| --- | --- | --- | --- | --- | --- |
|  | Percent Genes Filtered Out | | | | |
| Dataset | None | 0.1 | 1 | 5 | 10 |
| Biase | 0.861 | 0.795 | 0.795 | **1.000** | **1.000** |
| Baron_human1 | **0.512** | 0.272 | 0.208 | 0.078 | 0.078 |
| Baron_human2 | **0.301** | 0.299 | 0.252 | 0.132 | 0.028 |
| Baron_human3 | 0.556 | 0.530 | 0.341 | 0.565 | **0.784** |
| Baron_human4 | **0.639** | 0.618 | 0.441 | 0.555 | 0.562 |
| Baron_mouse1 | **0.395** | 0.140 | 0.119 | 0.121 | 0.034 |
| Darmanis | **0.898** | **0.898** | 0.825 | 0.834 | 0.874 |
| Deng | 0.440 | 0.416 | 0.399 | 0.461 | **0.465** |
| Goolam | 0.575 | 0.670 | 0.615 | **0.933** | 0.424 |
| Li | 0.659 | 0.634 | **0.959** | 0.929 | 0.943 |
| Ting | 0.625 | 0.625 | 0.619 | **0.749** | 0.601 |
| Yan | **0.791** | **0.791** | **0.791** | 0.623 | 0.685 |
| Zeisel | 0.420 | 0.373 | 0.327 | **0.469** | 0.462 |

|  | ARI for t-SNE+*k-*means clustering | | | | |
| --- | --- | --- | --- | --- | --- |
|  | Percent Genes Filtered Out | | | | |
| Dataset | None | 0.1 | 1 | 5 | 10 |
| Biase | **0.948** | 0.695 | 0.695 | **0.948** | **0.948** |
| Baron_human1 | 0.512 | 0.431 | **0.532** | 0.433 | 0.514 |
| Baron_human2 | 0.842 | 0.839 | **0.877** | 0.765 | 0.871 |
| Baron_human3 | 0.623 | **0.897** | 0.894 | 0.866 | 0.857 |
| Baron_human4 | 0.745 | 0.730 | **0.814** | 0.757 | 0.738 |
| Baron_mouse1 | 0.593 | **0.620** | 0.586 | 0.598 | 0.592 |
| Darmanis | **0.792** | **0.792** | **0.792** | **0.792** | **0.792** |
| Deng | 0.438 | 0.438 | 0.438 | 0.430 | **0.443** |
| Goolam | **0.875** | 0.543 | 0.646 | 0.630 | 0.768 |
| Li | **0.903** | 0.897 | **0.903** | **0.903** | **0.903** |
| Ting | 0.854 | **0.894** | 0.866 | 0.803 | 0.723 |
| Yan | **0.621** | **0.621** | **0.621** | **0.621** | **0.621** |
| Zeisel | **0.782** | 0.775 | 0.776 | 0.780 | 0.770 |

|  | ARI for SIMLR clustering | | | | |
| --- | --- | --- | --- | --- | --- |
|  | Percent Genes Filtered Out | | | | |
| Dataset | None | 0.1 | 1 | 5 | 10 |
| Biase | 0.583 | **1.000** | **1.000** | **1.000** | **1.000** |
| Baron_human1 | **0.575** | 0.533 | 0.503 | 0.280 | 0.515 |
| Baron_human2 | 0.752 | 0.804 | 0.806 | **0.839** | 0.737 |
| Baron_human3 | **0.801** | 0.764 | 0.741 | 0.788 | 0.733 |
| Baron_human4 | 0.578 | 0.505 | 0.552 | **0.881** | 0.674 |
| Baron_mouse1 | **0.663** | 0.507 | 0.645 | 0.488 | 0.281 |
| Darmanis | 0.543 | 0.593 | **0.637** | 0.450 | 0.619 |
| Deng | 0.235 | 0.235 | 0.235 | 0.136 | **0.418** |
| Goolam | **0.471** | 0.382 | 0.394 | 0.389 | 0.389 |
| Li | 0.845 | 0.845 | **0.952** | 0.893 | 0.956 |
| Ting | **0.788** | **0.788** | **0.788** | **0.788** | 0.767 |
| Yan | **0.492** | **0.492** | **0.492** | 0.410 | 0.410 |
| Zeisel | 0.584 | 0.571 | 0.556 | 0.684 | **0.690** |

**Supplementary Table 7. Difference in ARI for CIDR, t-SNE+*k-*means, SIMLR, and SAME-clustering, with and without gene filtering.** Negative values indicate that gene filtering step led to improved performance. Last row (sd) shows the standard deviation of these differences across datasets.

| Dataset | CIDR | t-SNE+*k-*means | SIMLR | SAME |
| --- | --- | --- | --- | --- |
| Biase | -0.139 | 0.000 | -0.417 | -0.052 |
| Darmanis | 0.023 | 0.000 | -0.076 | 0.013 |
| Deng | -0.025 | -0.005 | -0.182 | -0.051 |
| Goolam | 0.152 | 0.045 | 0.082 | -0.320 |
| Li | -0.284 | 0.000 | -0.111 | 0.017 |
| Ting | 0.023 | 0.139 | 0.021 | -0.002 |
| Yan | 0.106 | 0.000 | 0.162 | 0.000 |
| Zeisel | -0.042 | 0.012 | -0.106 | 0.011 |
| hum1 | 0.434 | 0.003 | 0.060 | 0.096 |
| hum2 | 0.273 | -0.032 | 0.015 | 0.175 |
| hum3 | -0.228 | 0.034 | 0.067 | 0.064 |
| hum4 | 0.077 | 0.012 | -0.096 | -0.221 |
| mouse1 | 0.361 | 0.000 | 0.382 | -0.106 |
| mouse2 | 0.437 | -0.041 | 0.021 | -0.123 |
| sd | 0.229 | 0.043 | 0.182 | 0.127 |

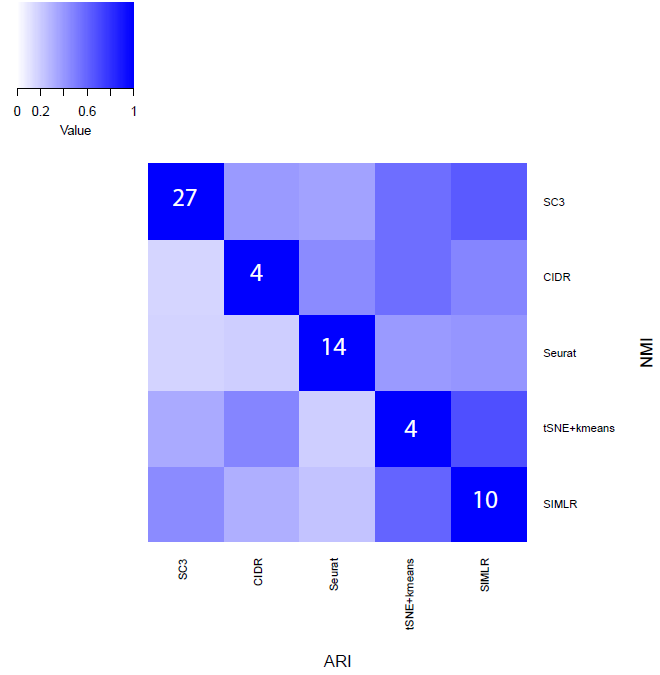

**Supplementary Figure 1. Similarity of clustering solutions from different methods.**

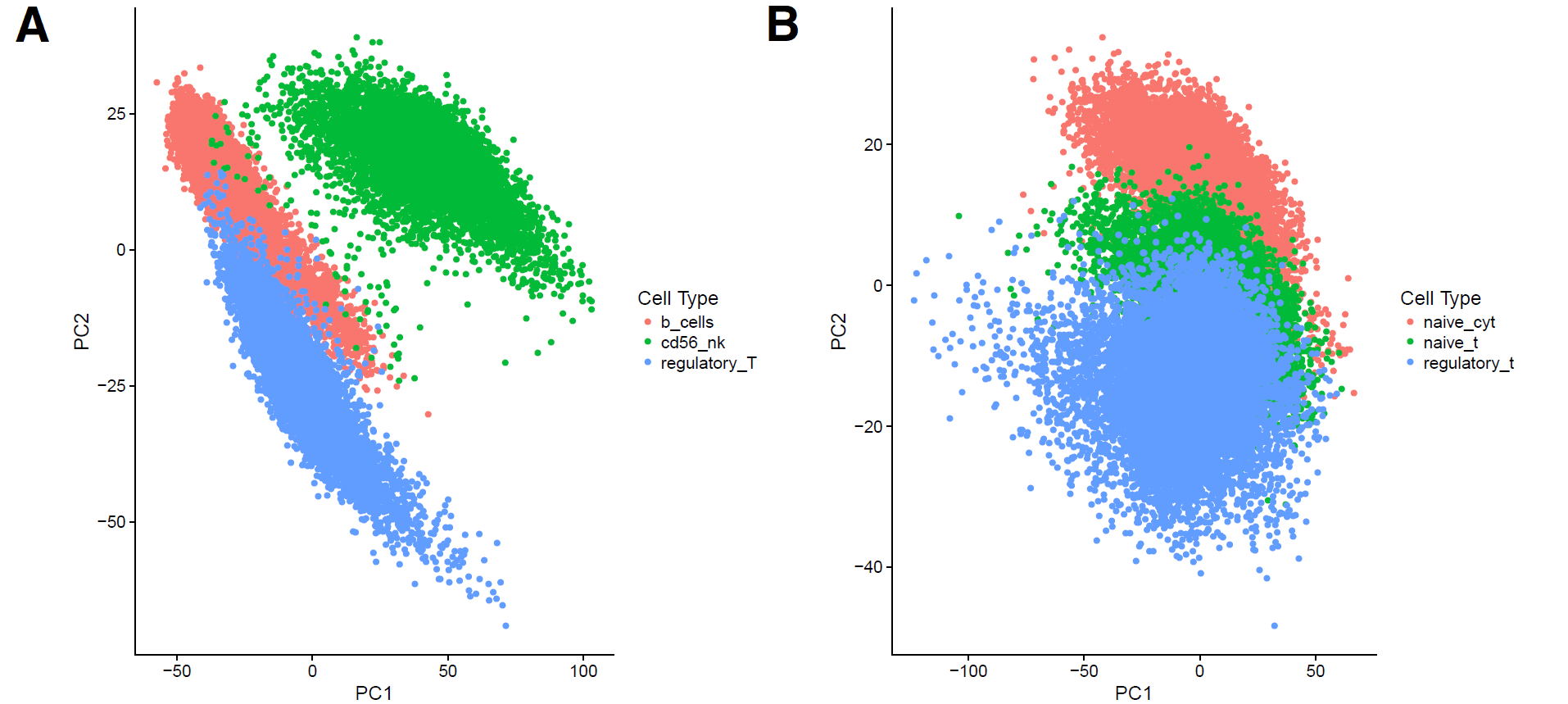

**Supplementary Figure 2. Visual inspection to determine number of clusters for simple and challenging case.** PCA plots of simple (A) and challenging (B) case using the top 1000 variable genes, based on log_2_ transformed and normalized expression matrix.

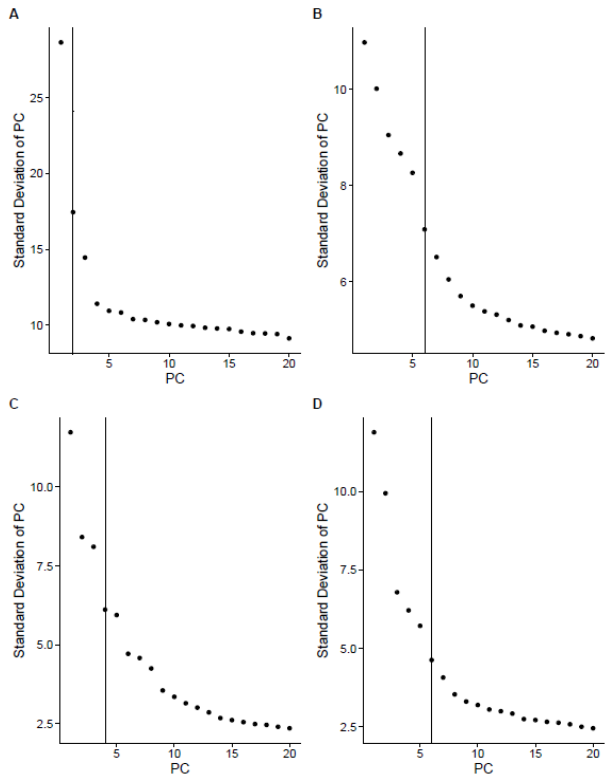

**Supplementary Figure 3. Elbow Plot.** Standard deviation of PCs is plotted to determine number of PCs to use for Seurat clustering. A-D correspond to Biase, Darmanis, Baron_Human1, and Baron_Human4 datasets, respectively. Line indicates the number of PCs selected by *nPC* function of CIDR.

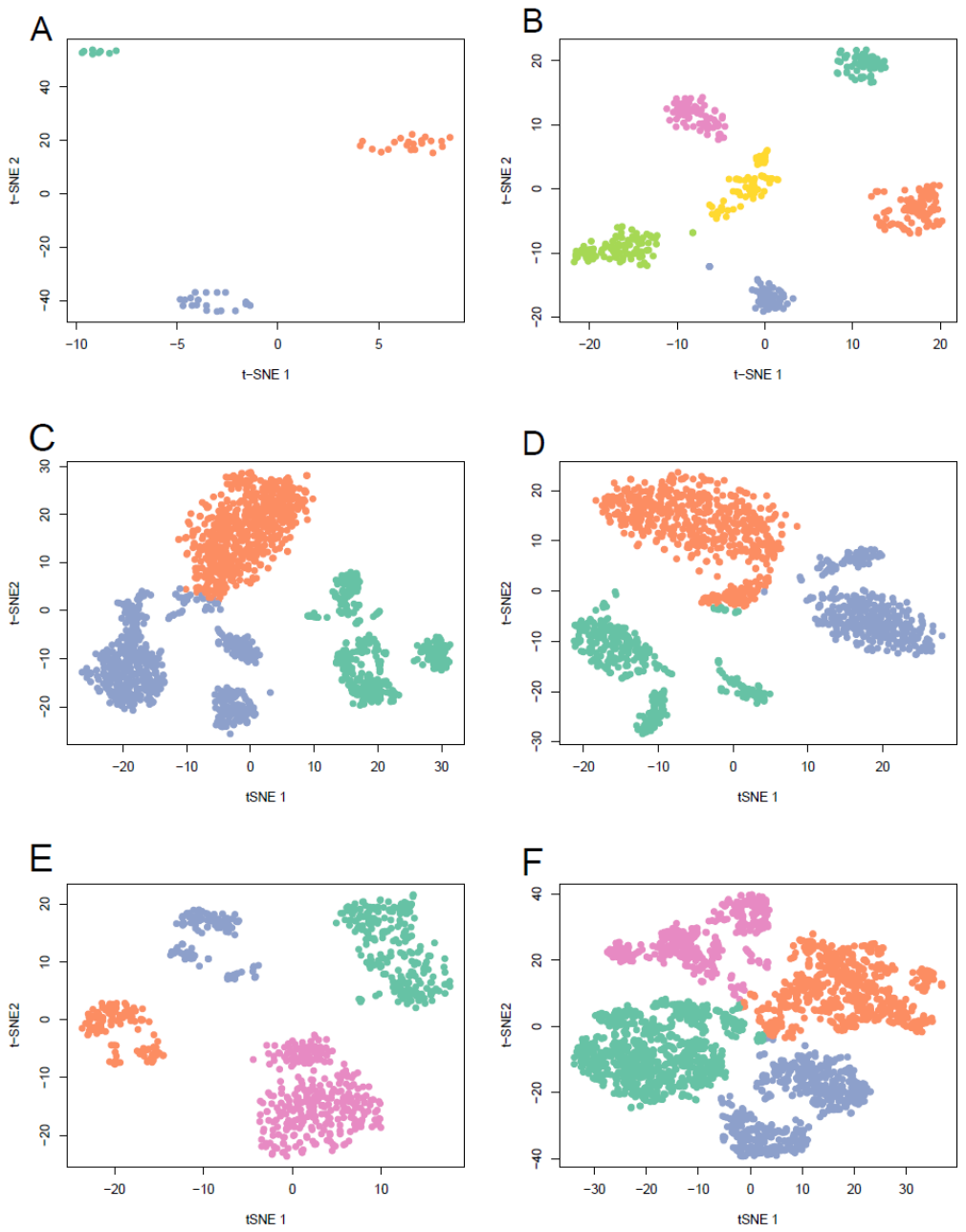

**Supplementary Figure 4. Visualization of single cell data using t-SNE.** t-SNE plot for t-SNE+*k-*means clustering with centroids determined automatically from ADPclust for Biase, Li, Baron_hum2, Baron_hum4, Baron_mouse1, and Zeisel datasets respectively. Colors in graph reflect t-SNE+*k-*means clustering labels.

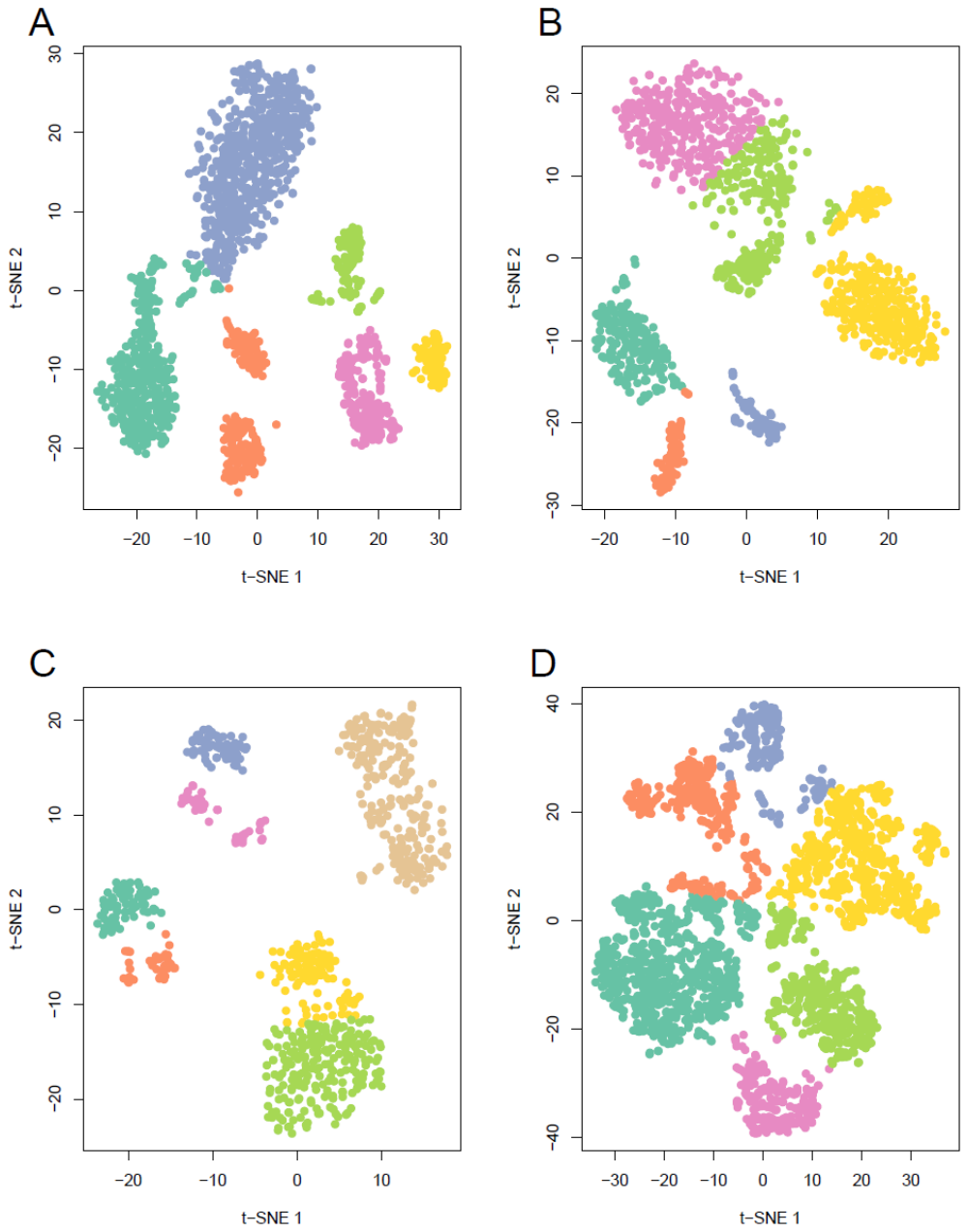

**Supplementary Figure 5. Adjusted visualization of single cell data using t-SNE.** t-SNE plot of t-SNE+*k-*means clustering results with manually selected centroids (based on visual inspection) for Baron_hum2, Baron_hum4, Baron_mouse1, and Zeisel datasets respectively. Colors in graph reflect t-SNE+*k-*means clustering labels.

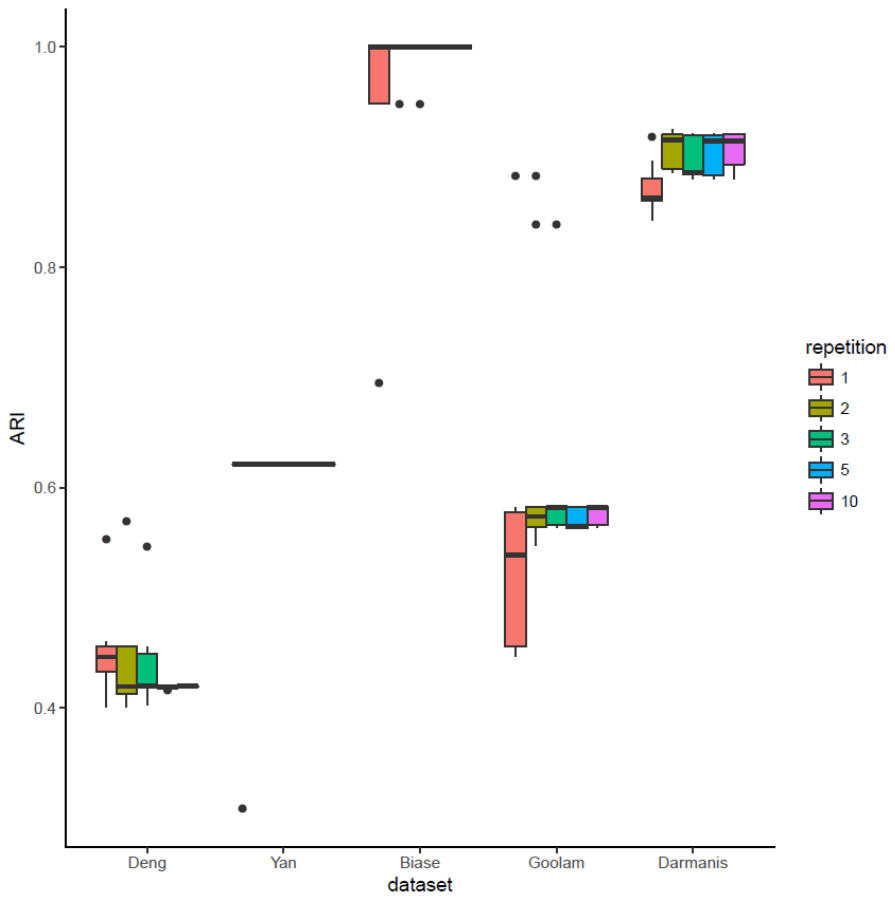

**Supplementary Figure 6. Number of EM Chains Needed to Stabilize ARI.** Variation in ARI for 5 datasets when EM Algorithm is run 1, 2, 3, 5, and 10 times, each retaining the one with maximum likelihood.

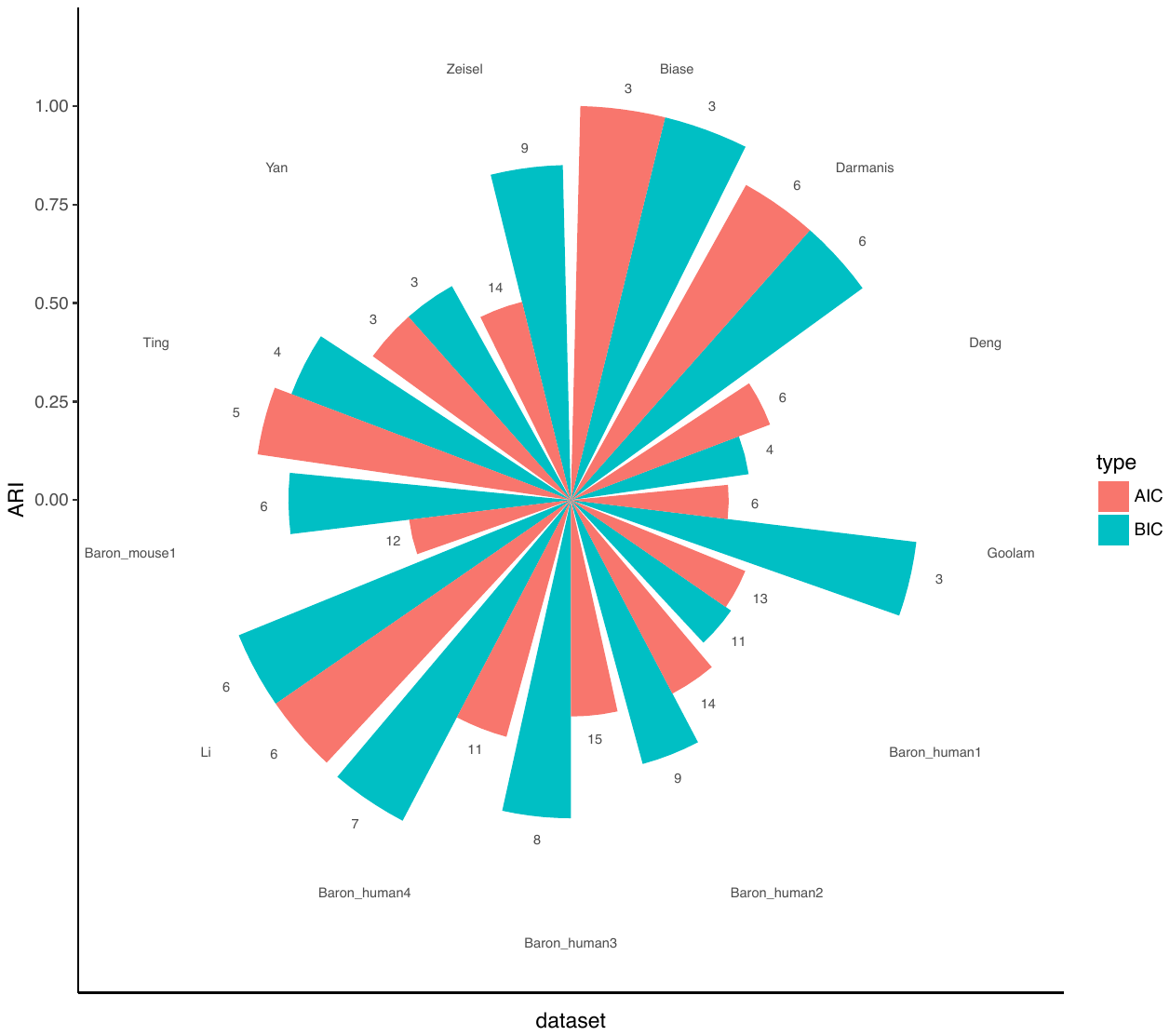

**Supplementary Figure 7. AIC vs BIC.** ARI across datasets where the estimated number of clusters is chosen by either AIC or BIC.

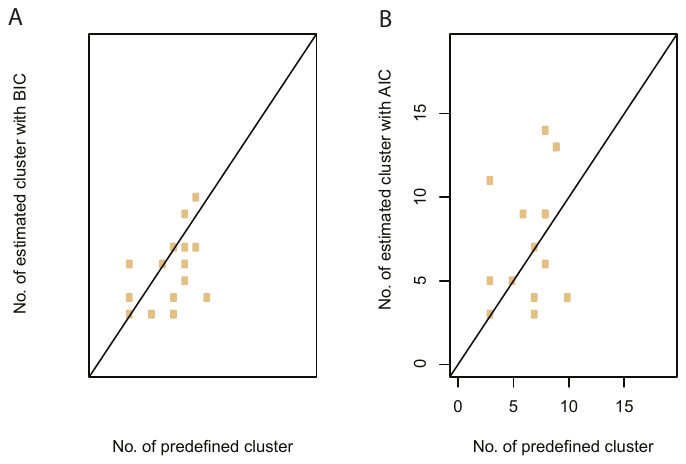

**Supplementary Figure 8. Correlations between the cluster numbers inferred by BIC (A) and AIC (B), respectively, and the true cluster numbers across the 15 benchmarking datasets.**

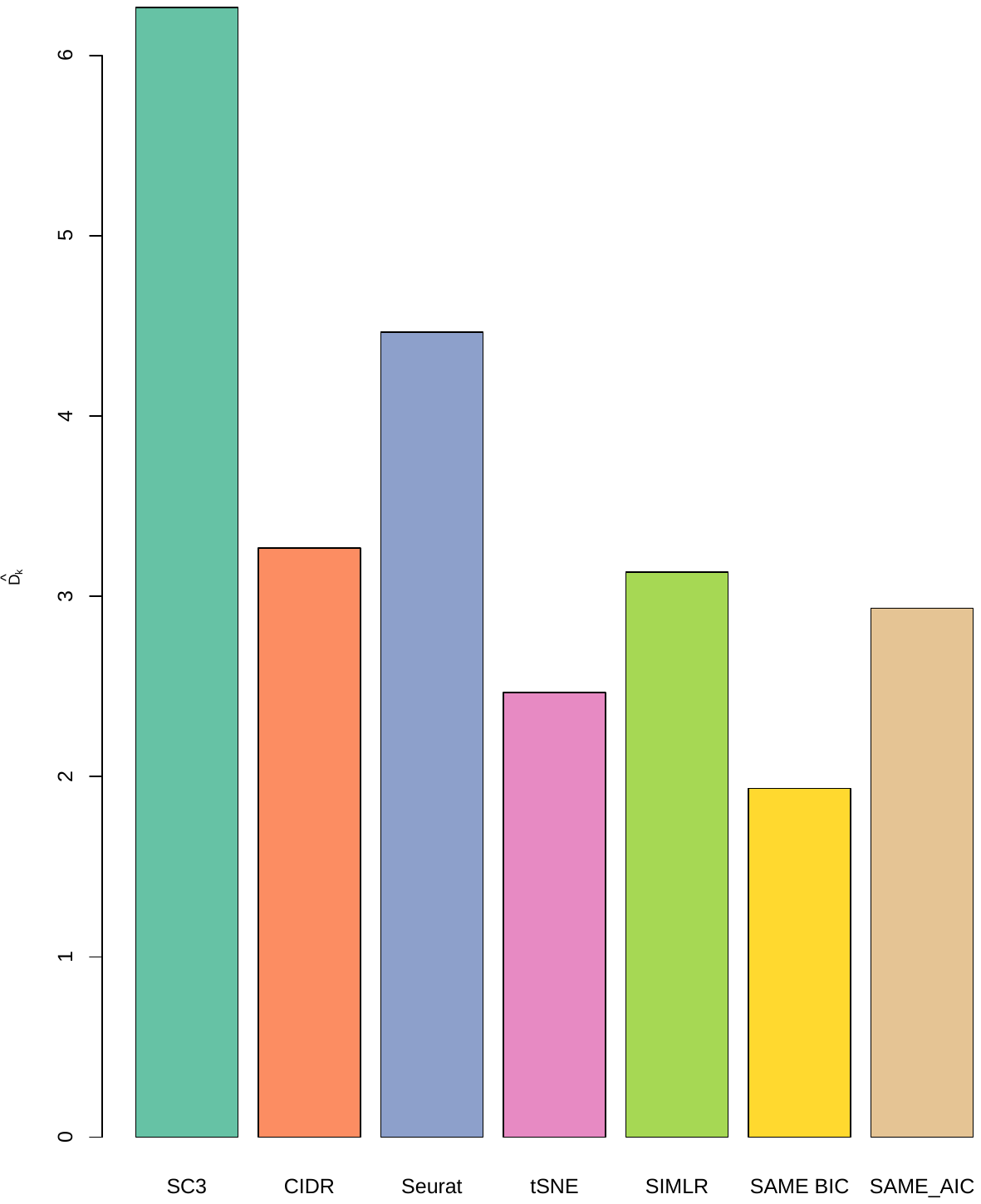

**Supplementary Figure 9. Average deviations between the cluster numbers inferred by five individual methods and SAME-clustering using BIC and AIC criteria, respectively, and the true cluster numbers, measured by** $\bar{\boldsymbol{D}}\boldsymbol{=}\frac{\boldsymbol{1}}{\boldsymbol{m}}\sum_{\boldsymbol{m}} \boldsymbol{|}\hat{\boldsymbol{k}}\boldsymbol{-}\boldsymbol{k}_{\boldsymbol{t}}\boldsymbol{|}$**, where the number of datasets m equals to 15.**

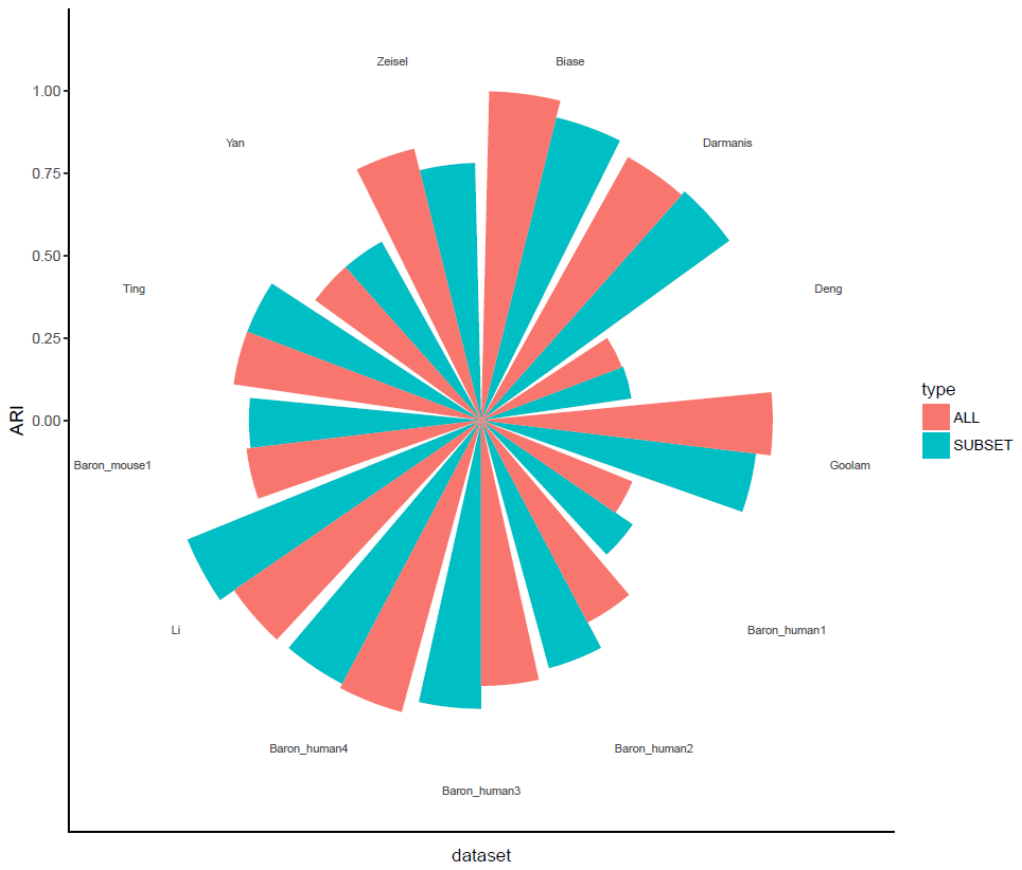

**Supplementary Figure 10.** **Ensemble of all five methods vs ensemble of four maximally diverse methods.** ARIs from two ensemble strategies, including either all five individual clustering solutions or the most diverse set of four solution, are compared.

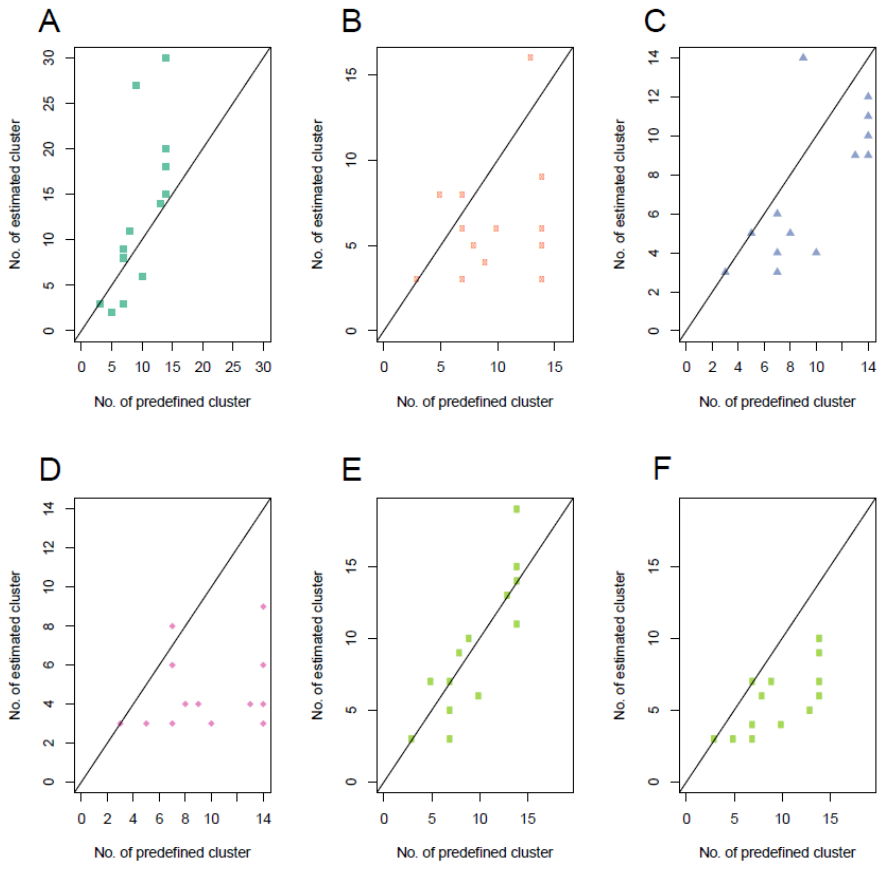

**Supplementary Figure 11. Accuracy of estimated number of clusters**. A-E. Correlations between estimated and true numbers of clusters across 13 benchmark datasets, for SC3 (A), CIDR (B), Seurat (C), t-SNE+*k-*means (D), SIMLR (E) and SAME-clustering (F).

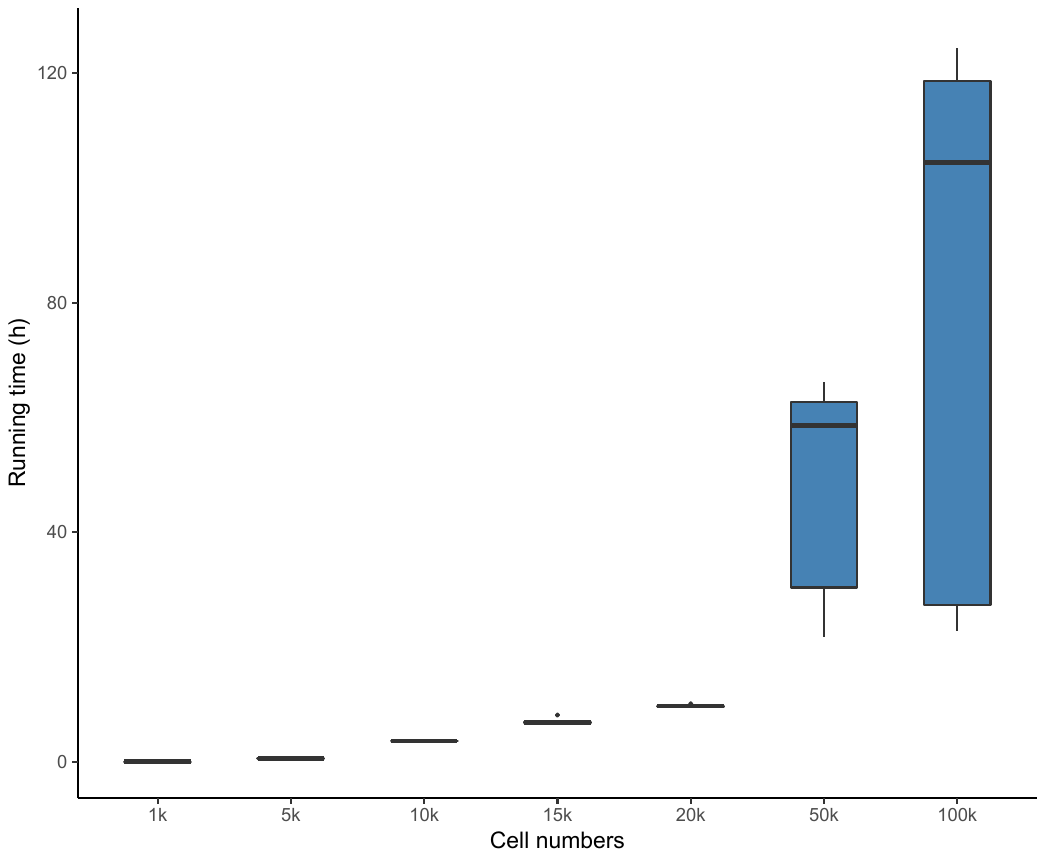

**Supplementary Figure 12. Running time of SAME-clustering for the PBMC mixture datasets with 1,000 - 100,000 single cell. Running time (Y-axis) is in hours (h).**

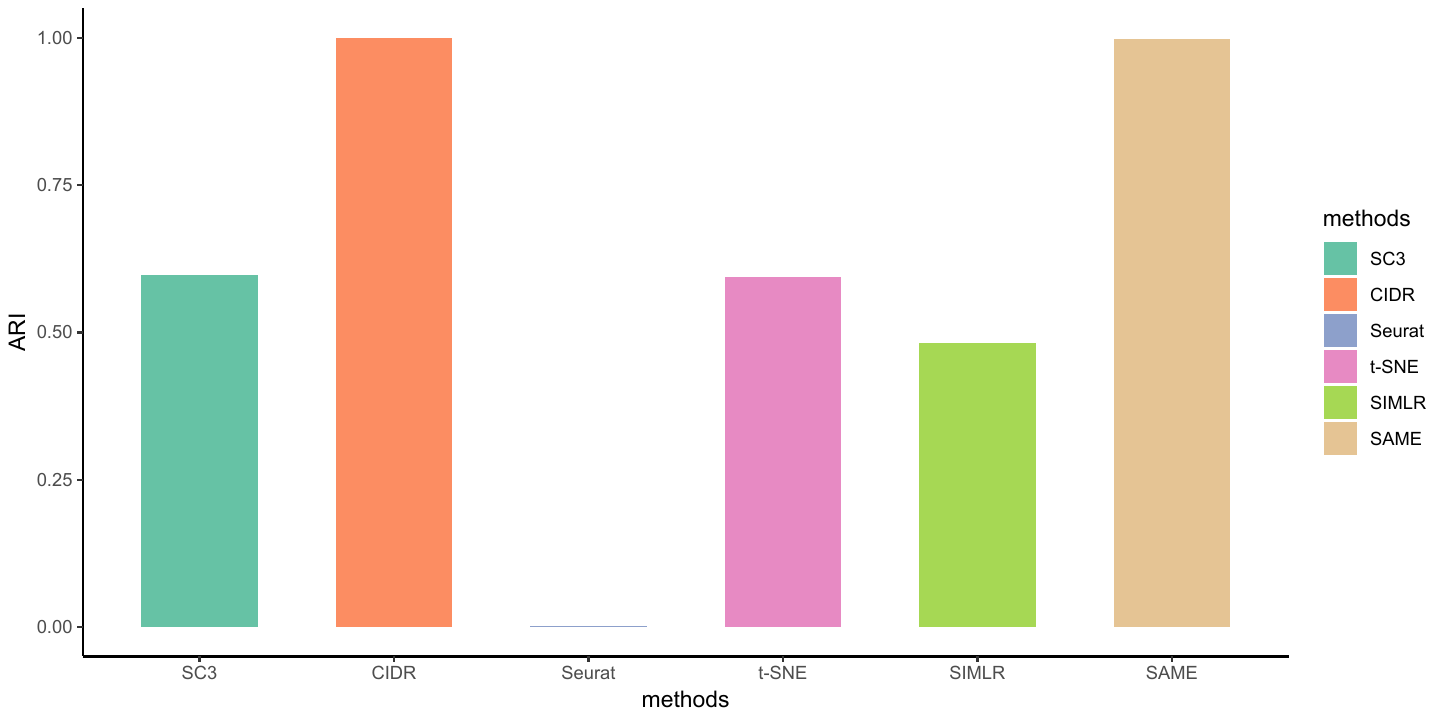

**Supplementary Figure 13. Benchmarking of SAME-clustering in simulated dataset.**

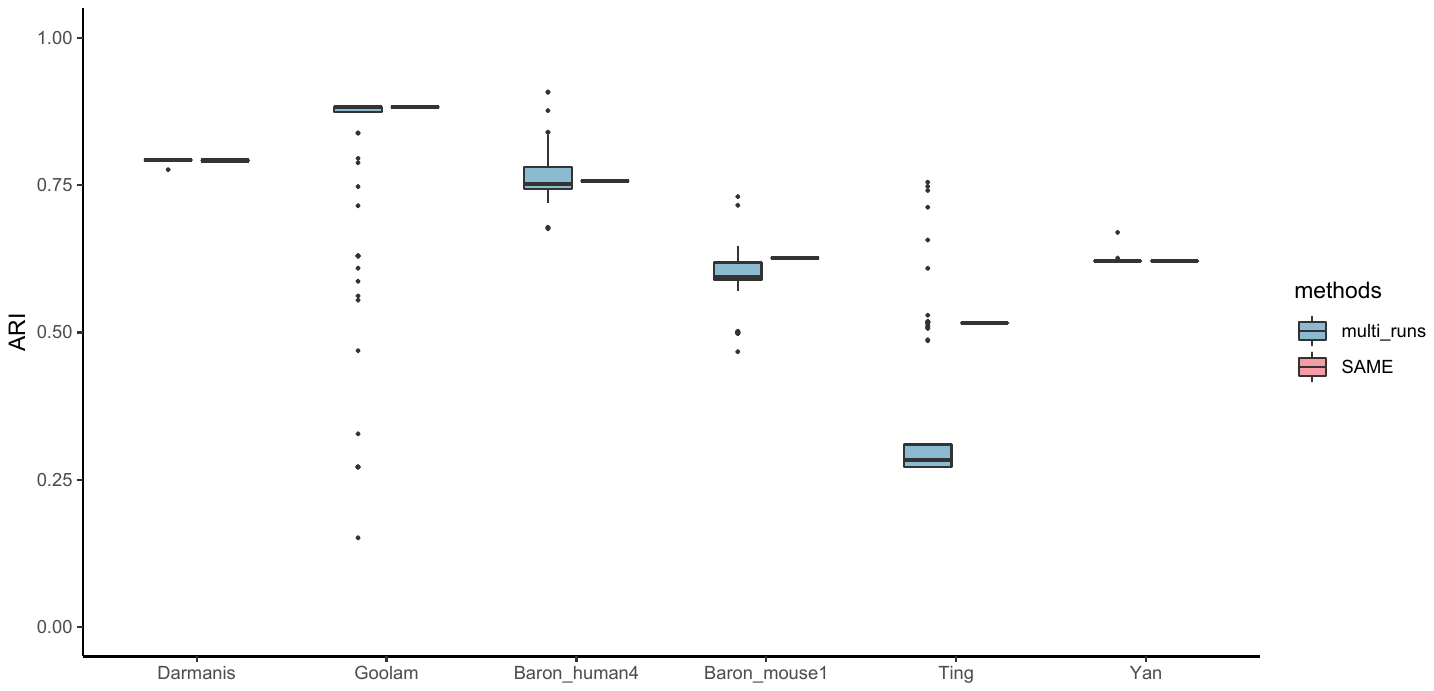

**Supplementary Figure 14. Evaluation of SAME’s robustness to multiple runs of one original individual clustering algorithm.** We first applied t-SNE + k-means method to six datasets for 100 times with different seeds (blue, left set of box plots), and then performed cluster ensemble using SAME-clustering (red, right set of box plots).

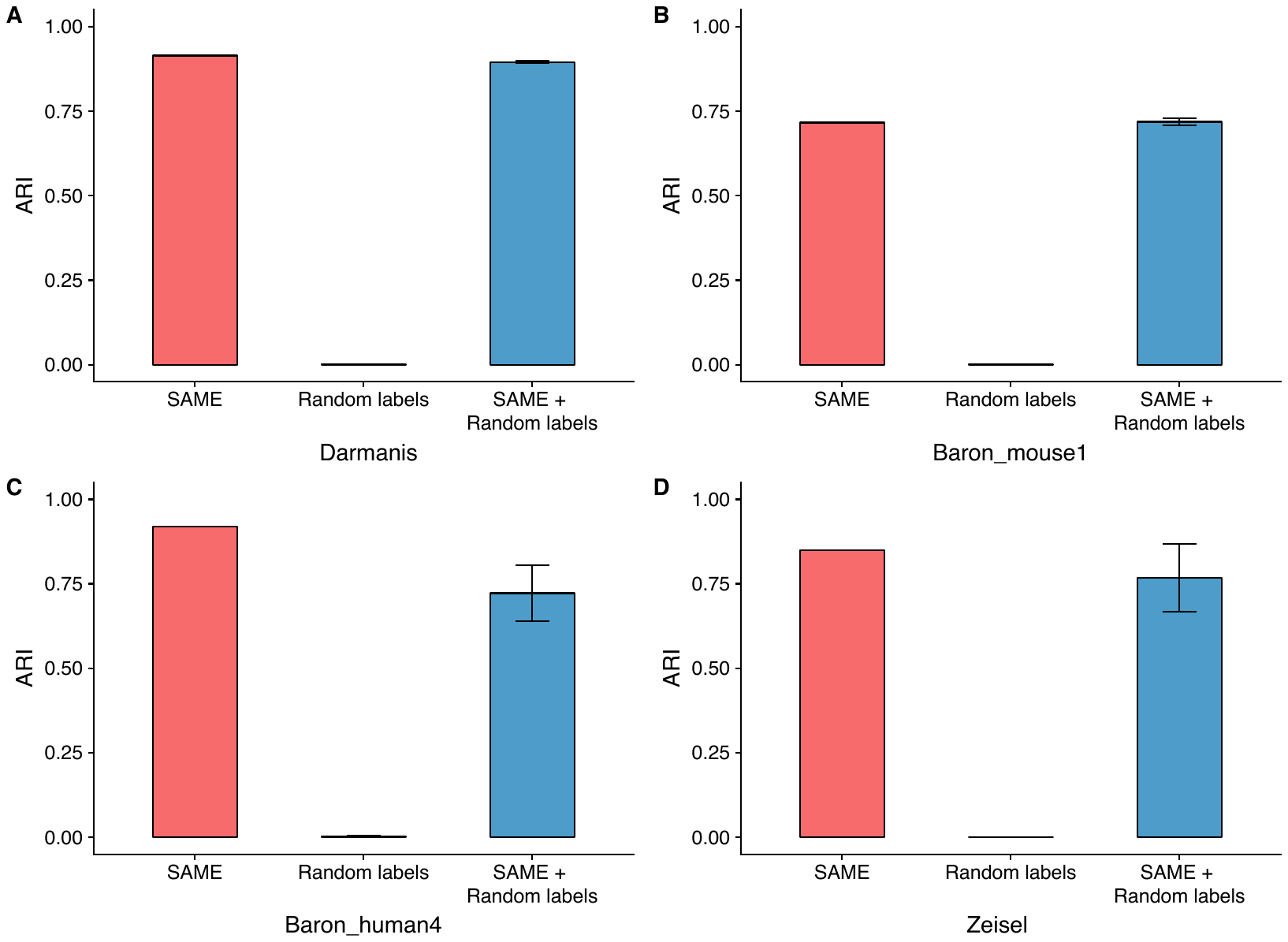
**Supplementary Figure 15. Evaluation of SAME’s robustness to random cluster labels.**

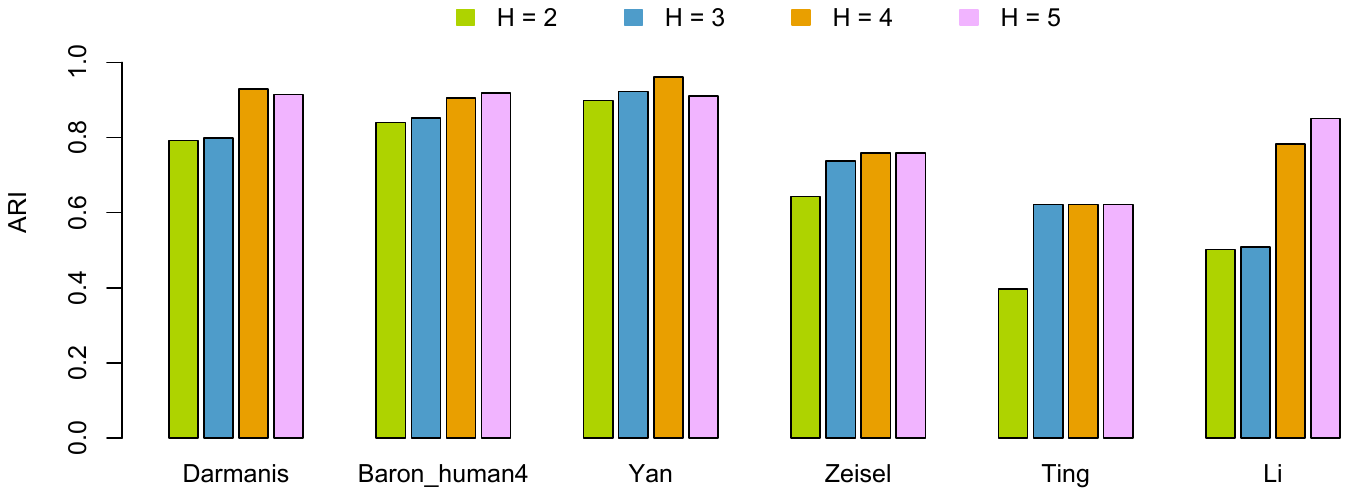

**Supplementary Figure 16. Impact of different numbers of input solutions (H = 2, 3, 4, and 5) on SAME’s performance.**

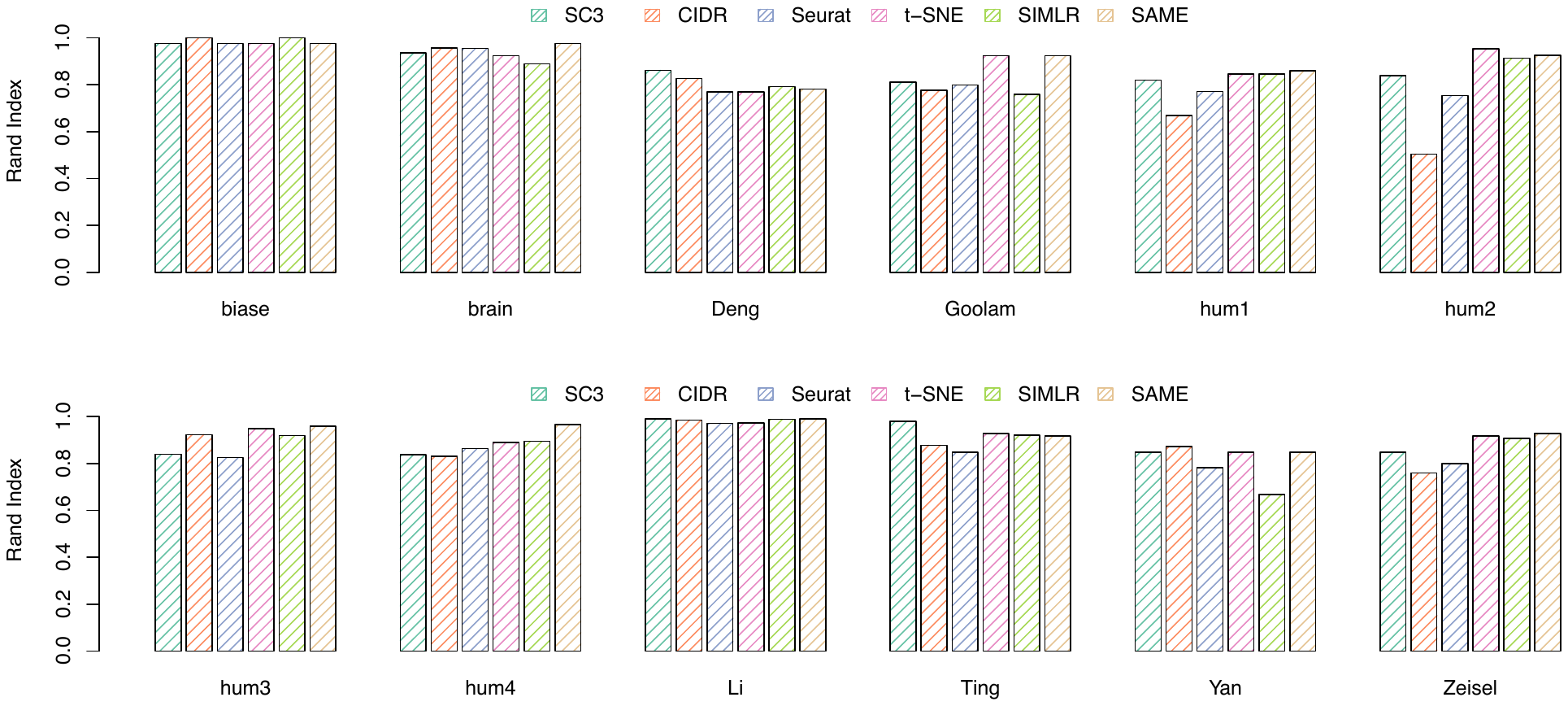

**Supplementary Figure 17. Benchmarking of SAME-clustering in 12 published datasets, using Rand Index to measure the similarity between inferred and true cluster labels.**

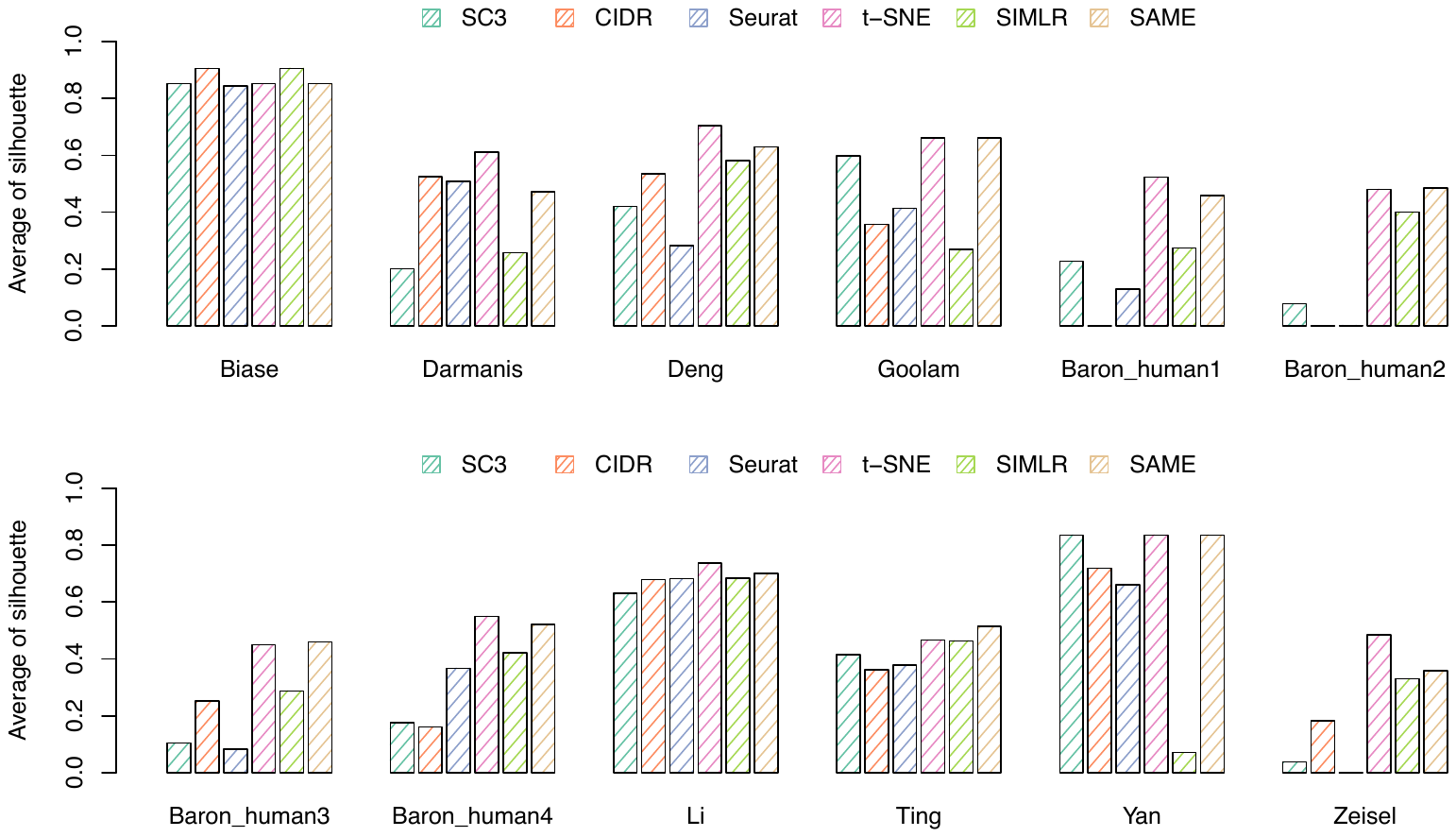

**Supplementary Figure 18. Benchmarking of SAME-clustering in 12 published datasets, using silhouette width to measure the cluster tightness.**

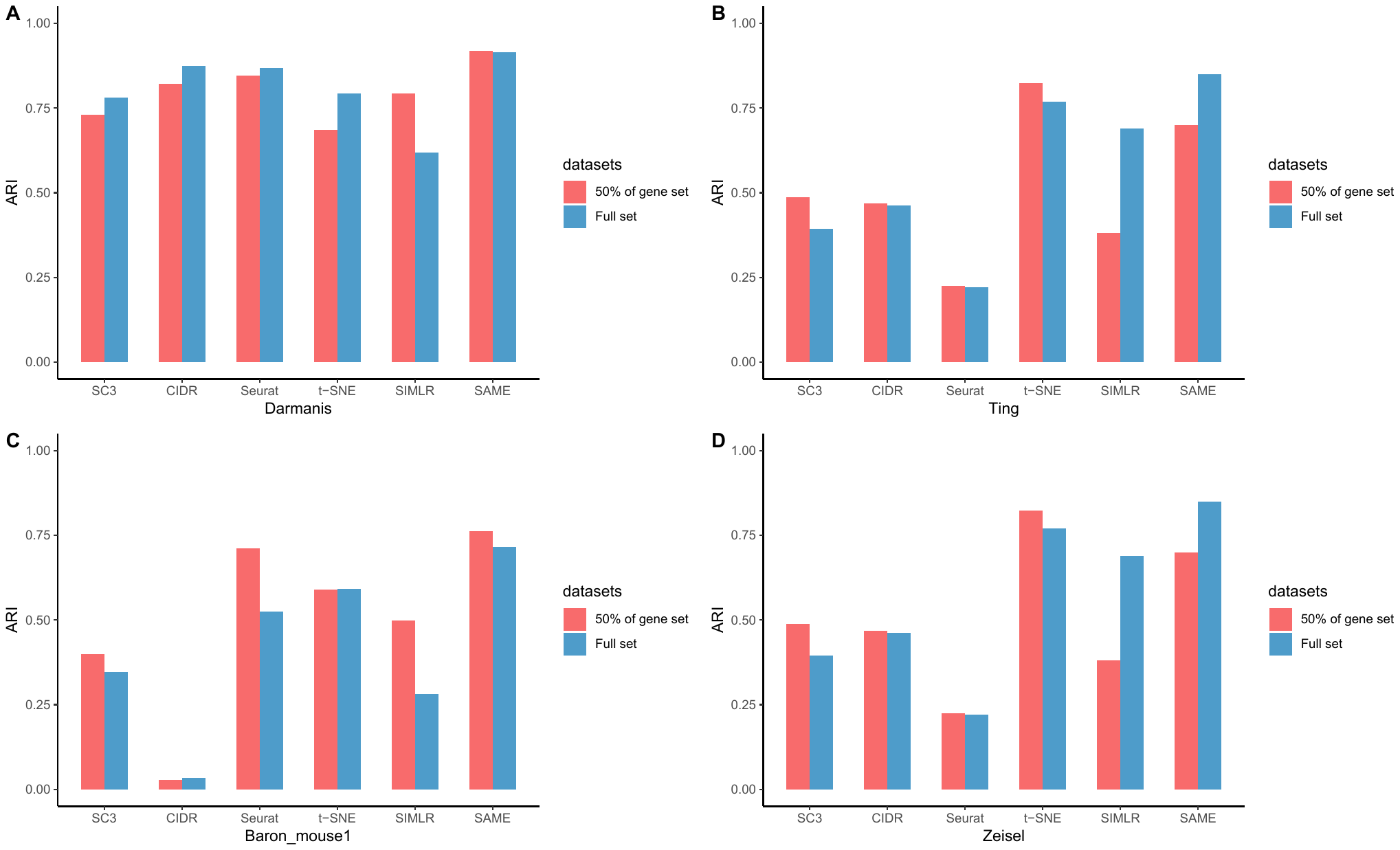

**Supplementary Figure 19. Evaluation the influence of different gene sets used on clustering accuracy of SAME-clustering.**

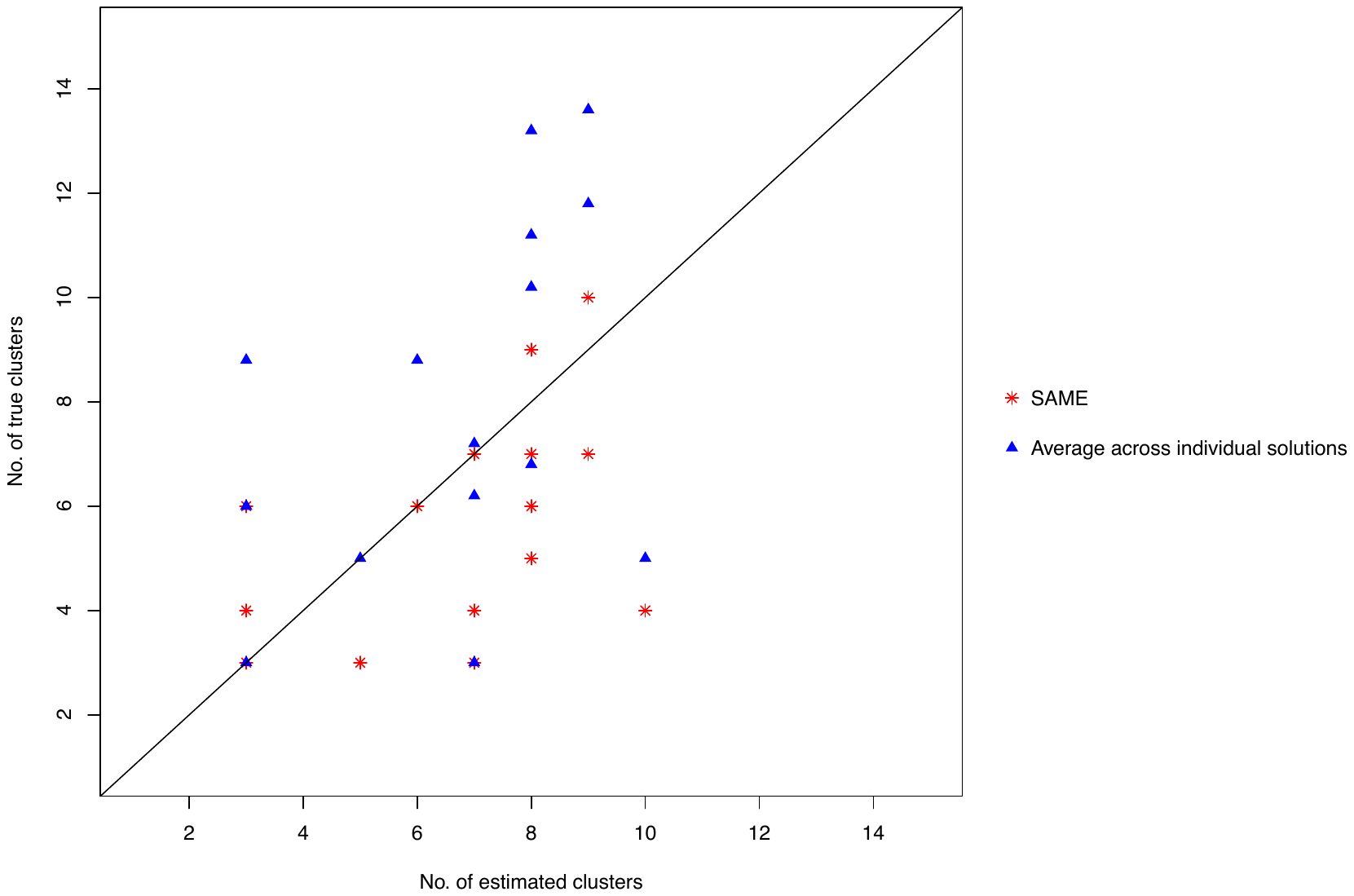

**Supplementary Figure 20. Average deviations between the inferred and the true numbers of clusters, measured by** $\bar{\boldsymbol{D}}\boldsymbol{=}\frac{\boldsymbol{1}}{\boldsymbol{m}}\sum_{\boldsymbol{m}} \boldsymbol{|}\hat{\boldsymbol{k}}\boldsymbol{-}\boldsymbol{k}_{\boldsymbol{t}}\boldsymbol{|}$**, where the number of datasets m equals to 15.**

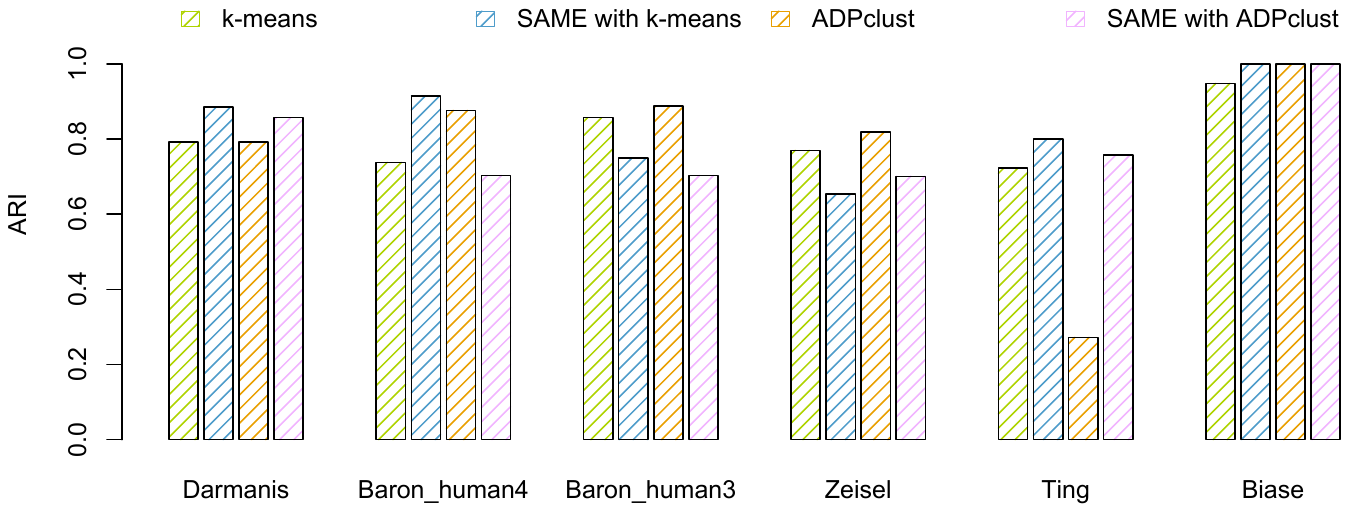

**Supplementary Figure 21. Comparing performance of t-SNE + ADPclust and t-SNE + ADPclust + k-means in six datasets (Darmanis, Baron_human4, Baron_human3, Zeisel, Ting and Biase).**

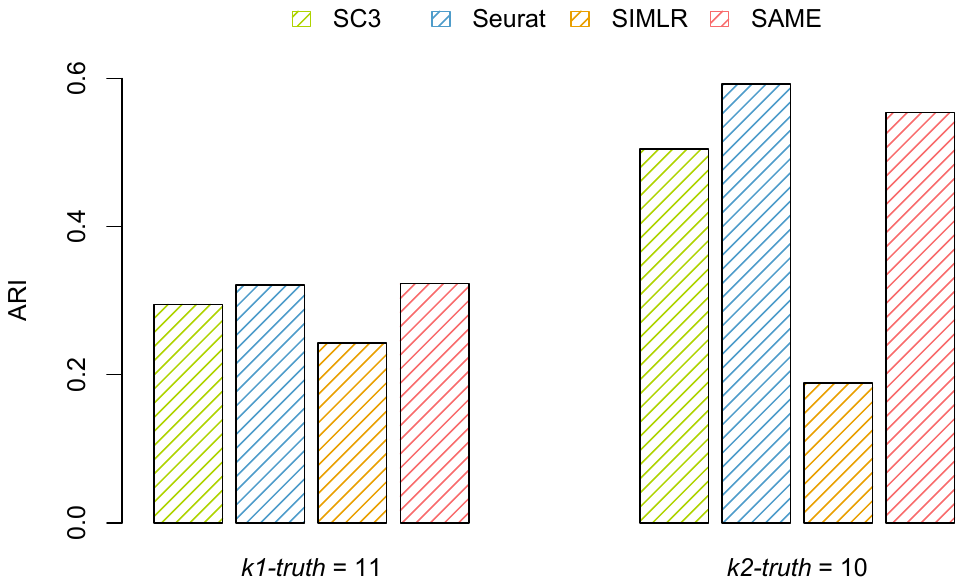

**Supplementary Figure 22. Evaluation the SAME’s performance in large and complex “Fresh 68k” PBMC dataset.**

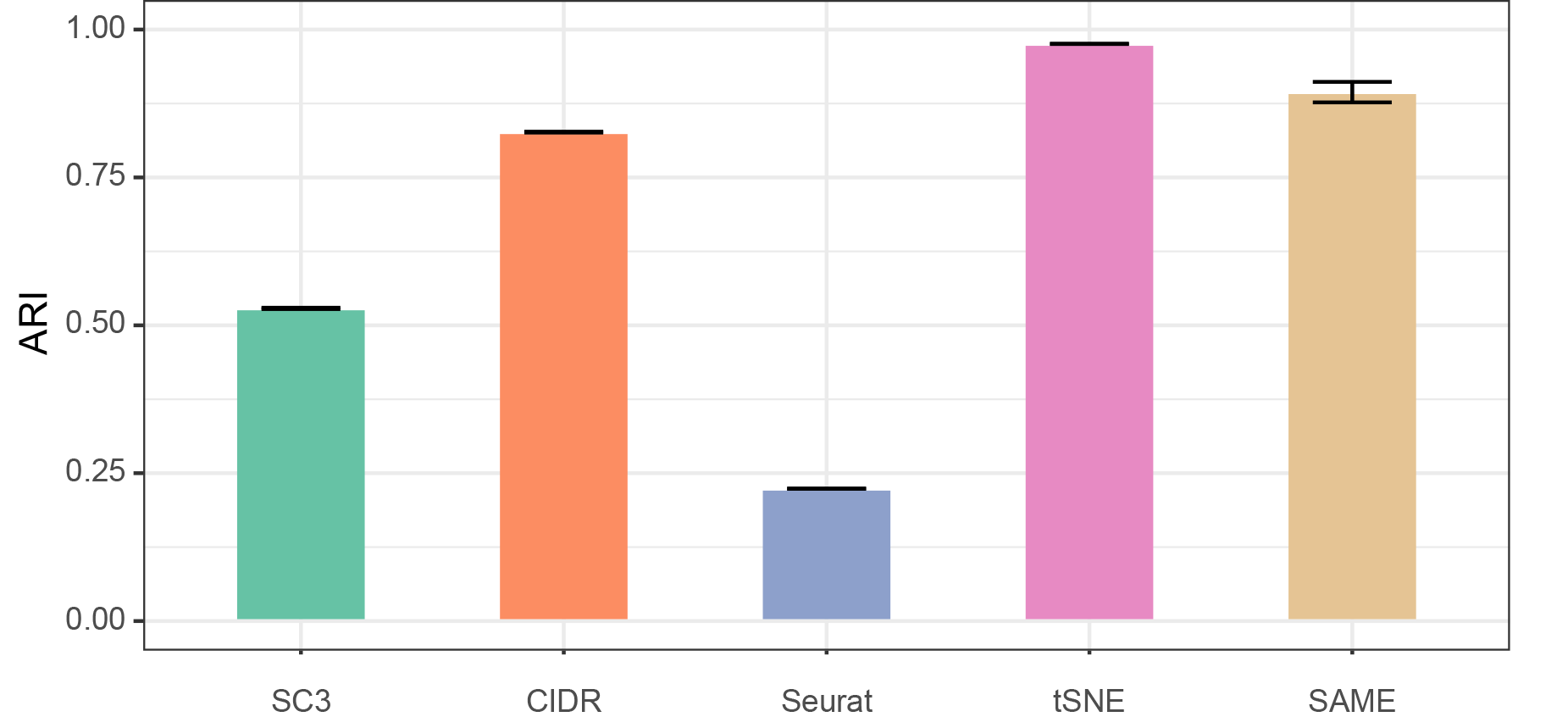

**Supplementary Figure 23. Evaluation the SAME’s performance in a simulated large dataset of and 100k single cells.**

**Software versions used for individual methods**

R package versions used are SC3 1.7.2, Seurat 2.1.0. Cidr 0.1.5, Rtsne 0.13, ADPclust 0.7, and SIMLR 1.4.1. SAME-clustering if freely available at <https://yunliweb.its.unc.edu/same/> and <https://github.com/yycunc/SAMEclustering>.
